# Inositol phosphates, pyrophosphates and the genes involved in their turnover in the streptophyte green alga *Chara braunii*

**DOI:** 10.64898/2026.03.30.715254

**Authors:** Daniel A. Heß, Anuj Shukla, Henning Jacob Jessen, Wolfgang R. Hess

**Affiliations:** Genetics and Experimental Bioinformatics Group, Faculty of Biology, University of Freiburg, Schänzlestr. 1, 79104 Freiburg, Germany; Institute of Organic Chemistry, University of Freiburg, Albertstr. 21l, D-79104 Freiburg, Germany; CIBSS-Centre for Integrative Biological Signalling Studies, University of Freiburg, Germany

## Abstract

Inositol phosphates (InsPs) and inositol pyrophosphates (PP-InsPs) are conserved signalling molecules, but their evolutionary origin and diversification in the green lineage remain poorly understood. Here we investigated the InsP network in the streptophyte alga *Chara braunii*, a key lineage close to the origin of land plants. Using capillary electrophoresis–electrospray ionization mass spectrometry, we detected a broad spectrum of InsP and PP-InsP species from InsP_3_ to InsP_8_, including multiple positional isomers. Developmental profiling across dormant oospores, young thalli and mature thalli revealed extensive metabolic remodeling, with InsP_6_ as the dominant metabolite and distinct stage-dependent changes in lower InsPs and pyrophosphorylated species. Multiple PP-InsP_5_ and (PP)_2_-InsP_4_ isomers were identified, together with an unassigned additional InsP_8_-like signal, indicating further pathway complexity. Bioinformatic analyses identified candidate homologs of major InsP metabolic enzymes, supporting the presence of an enzymatic framework for InsP synthesis and turnover similar to land plants. Environmental perturbation revealed isomer-selective effects: prolonged light and dark phases strongly affected the accumulation of specific InsP_5_ and PP-InsP_5_ isomers, with 1-PP-InsP_5_ emerging as the most stimulus-responsive pyrophosphate species, whereas heat stress preferentially reduced 4-PP-InsP_5_. Together, these findings show that a structurally complex and environmentally responsive InsP network was already established in streptophyte algae before the emergence of land plants.

## Introduction

The macroscopic streptophyte alga *Chara braunii* is a prominent model in plant molecular and evolutionary developmental biology (Leebens-Mack et al., 2019; Kurtović et al., 2023) and together with *Nitella, Chara* is the most extensively studied genus within the class Charophyceae (McCourt et al., 1996; Foissner and Wasteneys, 2014). The Characeae are widespread and thrive in a variety of environmental conditions, from fresh to brackish waters, and they can withstand elevated salinity levels in nonmarine waters (Doege et al., 2016; McCourt et al., 2017). Different *Chara* species have served the scientific community for centuries (Corti, 1774), and gained significance as model species in (electro)physiological studies early on (Braun et al., 2007; Beilby and Casanova, 2014; Foissner and Wasteneys, 2014).

While the Charophyceae are not the direct sister group to land plants (Wickett et al., 2014; Cheng et al., 2019; Bierenbroodspot et al., 2024), they possess key ancestral traits shared with land plants and exhibit remarkable morphological complexity. They are the only class of streptophyte algae with rhizoids and tissue-like structures (Nishiyama et al., 2018; Bonnot et al., 2019; Fürst-Jansen et al., 2020). As a representative of the embryophyte algae precursors (Martin and Allen, 2018), *Chara* shares striking similarities, yet reveals also important differences in relation to land plants and therefore is a crucial model to examine within the Viridiplantae (Nishiyama et al., 2018).

Phosphorus is an essential nutrient, taken up primarily in the form of inorganic phosphate (Pi) by plants as well as by *Chara* (Andrews, 1987). Phosphorus is then utilized in molecules relevant for energy metabolism and signaling, among others in the form of inorganic polyphosphates (polyPs) (Sanz-Luque et al., 2020) and inositol phosphates (InsPs) (Irvine and Schell, 2001; Nagpal et al., 2024; Yang et al., 2024; Giehl and Schaaf, 2026). InsPs are found in all eukaryotes, where they play vital roles in various signal transduction pathways and regulate a broad spectrum of biological processes (Lorenzo-Orts et al., 2020; Riemer et al., 2022; Kim et al., 2024; Xiong et al., 2024). InsPs are composed of *myo*-inositol, a special cyclohexanehexol, to which different numbers of phosphate residues are esterified. A large array of InsP species can be generated *in vivo* through combinatorial phosphorylation, with further phosphorylation on existing phosphate groups (**Fig. 1**) resulting in inositol pyrophosphates (PP-InsPs). Some PP-InsPs can occur as mirror-images (enantiomers). In many cases, however, the exact enantiomer cannot yet be definitively assigned. To reflect this ambiguity, a notation such as 1/3-PP-InsP_5_ is used, indicating that the diphosphate group may be located at either the 1 or 3 position. In this publication, we report only one of the two possible isomers for simplicity, while acknowledging that the absolute stereochemistry remains unassigned.

**Figure 1.**
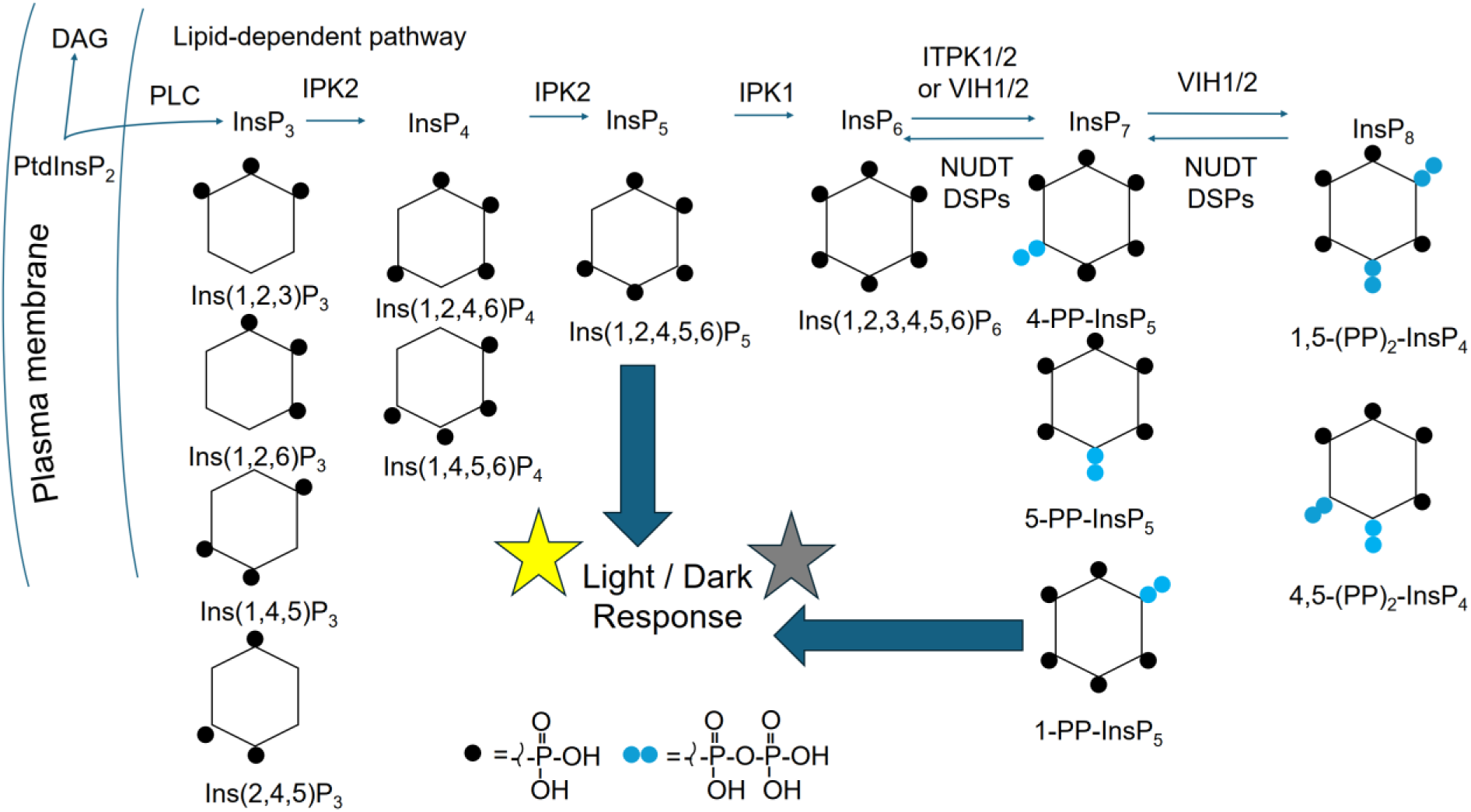
Simplified overview of the inositol pyrophosphate biosynthesis and metabolism in plants. Following either lipid-dependent or -independent routes of synthesis, inositol hexakisphosphate (InsP6) is phosphorylated by ITPK1/2 or VIH1/2 to generate 5-InsP_7_, 4/6-InsP_7_ and 1-InsP_7_, respectively. Detected isomers are indicated underneath the respective structures on top. 5-InsP_7_ can be further used as substrate by VIH1/2 and phosphorylated to generate 1,5-InsP_8_. PFA-DSPs and NUDTs may dephosphorylate the various inositol pyrophosphate species. Enzyme abbreviations: PLC, phospholipase C; IPK1, inositol-pentakisphosphate 2-kinase 1; IPK2, inositol polyphosphate 3-/6-/5-kinase 2; VIH, Inositol hexakisphosphate and diphosphoinositol-pentakisphosphate kinase, *Arabidopsis* homologs of yeast Vip; PFA-DSP, plant and fungi atypical dual-specificity phosphatases; NUDT, nudix hydrolase. The scheme is based on information from references (Williams et al., 2015; Lorenzo-Orts et al., 2020; Qiu et al., 2020; Ghosh et al., 2025) and it also incorporates the isomeric complexity detected in *Chara* during this study.

Both InsPs and PP-InsPs are present in all eukaryotes, though they vary in composition and concentration among different organisms and cell types (Shears, 2015; Lorenzo-Orts et al., 2020; Kim et al., 2024). However, the cellular targets and biosynthesis of these crucial messenger molecules have remained poorly understood throughout the Viridiplantae, even though first interactome maps are now becoming available (Sturm et al., 2025; Ritter et al., 2026).

Recent efforts have revealed both the conservation of crucial enzymes in plants (Yadav et al., 2025), as well as some striking differences in comparison to animals and fungi (Ghosh et al., 2025). In land plants, inositol phosphate metabolism follows a conserved kinase-driven pathway in which lower phosphorylated InsPs are sequentially converted into higher phosphorylated species by inositol polyphosphate multikinases (IPK2/IPMK) and inositol tris/tetrakisphosphate kinases (ITPKs), culminating in the formation of InsP_6_ by inositol pentakisphosphate 2-kinase (IPK1) (**Fig. 1**) (Kim et al., 2024; Giehl and Schaaf, 2026). In contrast to yeast and metazoans, plants lack canonical inositol hexakisphosphate kinases (IP6Ks) and instead use plant-specific ITPK1/ITPK2 enzymes to phosphorylate InsP_6_ and generate InsP_7_ isomers (Laha et al., 2019; Riemer et al., 2022), which are further converted into InsP_8_ by VIP-like (Mulugu et al., 2007) diphosphoinositol pentakisphosphate kinases (VIH) (**Fig. 1**) (Laha et al., 2015). These high-energy inositol pyrophosphates are now believed to function as central regulators of phosphate sensing (Dong et al., 2019; Zhu et al., 2019; Riemer et al., 2021). Importantly, the InsP network is dynamically remodelled under environmental stress conditions, including phosphate limitation, salt stress, heat, light fluctuations, and drought, where altered PP-InsP levels modulate SPX-domain signaling, transcriptional responses, and physiological acclimation (Laha et al., 2015; Wild et al., 2016; Couso et al., 2021; Shukla et al., 2021; Riemer et al., 2022; Ghosh et al., 2025).

In addition, PP-InsPs play crucial functions in other stress acclimation responses, often acting in signalling chains downstream of phytohormones like jasmonate, auxin and salicylic acid (Laha et al., 2015; Laha et al., 2016; Gulabani et al., 2022; Riemer et al., 2022). Signalling effects of PP-InsPs are conveyed through protein-protein interactions, allosteric regulation, non-enzymatic protein pyrophosphorylation and competition with phosphoinositides (Lorenzo-Orts et al., 2020; Nagpal et al., 2024; Lampe et al., 2025; Giehl and Schaaf, 2026). In green algae, InsP-related signalling is well documented in *Chlamydomonas* (Irvine et al., 1992; Couso et al., 2016; Bedera-García et al., 2025). However, there are no reports on the chemical composition of InsP species or on their differential accumulation under different growth conditions or the likely involved enzymes in multicellular green algae such as *Chara braunii*.

Because of its phylogenetic position at the transition between aquatic algae and land plants, the organization and regulation of the inositol phosphate network in the charophyte alga *Chara braunii* is of significant interest. Here, we combine metabolic profiling and bioinformatic analyses to investigate the architecture of InsP and PP-InsP metabolism in *Chara braunii*. Using capillary electrophoresis electrospray ionization mass spectrometry (CE-ESI-MS), a method that enables sensitive detection, quantitation, and isomer-resolved analysis of inositol phosphates, (Qiu et al., 2020; Liu et al., 2023; Qiu et al., 2023; Liu et al., 2025) we examine the presence and developmental dynamics of InsP_3_ to InsP_8_ species. We further reveal differential accumulation of PP-InsPs in *Chara braunii* under varying light and stress conditions. In parallel, comparative sequence analysis identifies genes encoding 14 candidate enzymes likely involved in InsP and PP-InsP biosynthesis. Together, these approaches establish how InsP metabolism operates in *Chara braunii* and provide insight into the evolutionary origin of plant inositol phosphate signaling and its role in environmental stress adaptation.

## Results

### Detection and developmental dynamics of inositol phosphate metabolism in Chara braunii

The charophyte alga *Chara braunii* represents an evolutionary intermediate between aquatic algae and land plants and provides a valuable system for investigating the origin and regulation of the InsP network in the green lineage. To study this network in *Chara braunii*, particularly how it is remodeled during development, CE-ESI-MS analysis was performed across three representative stages: dormant oospores, actively growing thalli at 12 days after germination (DAG), and mature thalli (40–60 days), see **Fig. 2**. Here, DAG refers to days after transfer following an initial 21-day germination period under standard growth conditions.

**Figure 2.**
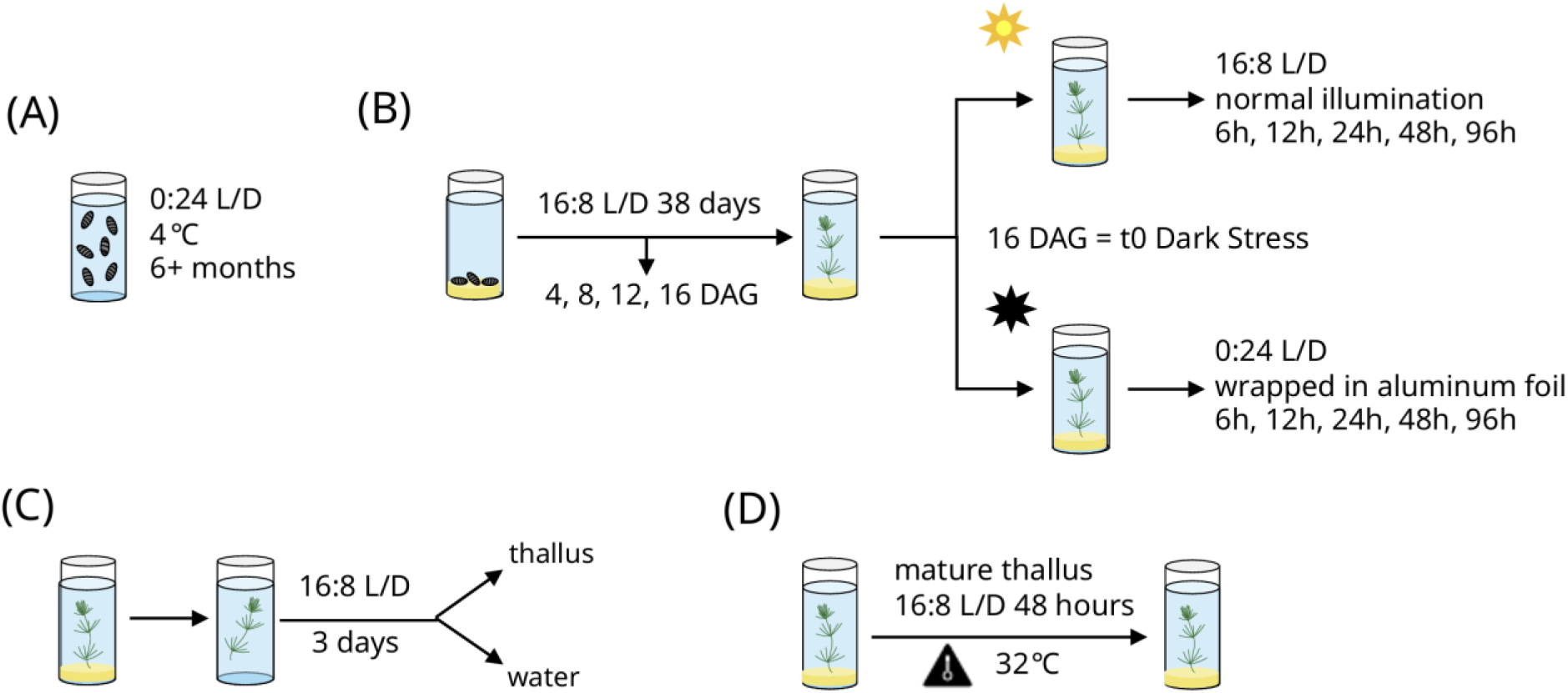
Schematic representation of the cultivation and experimental setup. **A)** Pooled, dormant oospores kept in darkness at 4°C for at least six months were collected. **B)** Thallus samples grown from oospores under standard conditions for a period of 38 days were harvested in triplicates. After an initial germination period of 21 days, samples were collected after four, eight, 12 and 16 days (DAG), with the last sample serving as t0 for the following Light/Dark incubation. Remaining culture vessels were divided into control (standard conditions) and dark treatment (wrapped in aluminium foil) during a regular mid-day light period. **C)** Triplicates of mature, vegetatively propagated thalli grown under standard conditions were harvested after three days in deionised water, or **D)** subjected to two days of 32°C heat stress.

Using TiO_2_ extraction (Wilson and Saiardi, 2018), we validated InsPs recovery from *C. braunii* extracts. Across development stages of *C. braunii*, a recovery rate of ∼60-65% was achieved (**Fig. S1A**). The obtained electropherograms revealed the presence of a broad spectrum of InsPs and PP-InsPs: including InsP_3_, InsP_4_, InsP_5_, InsP_6_, PP-InsP_4_ and further pyrophosphorylated derivatives (PP-InsP_5_ and (PP)_2_-InsP_4_), with clear separation of multiple positional isomers (for an example, see **Fig. 3A**). Notably, distinct isomeric profiles were observed for each InsP class. Because enantiomeric species cannot be resolved under our CE-ESI-MS conditions, isomer nomenclature was applied using subscript numbering. For example, the enantiomeric pair Ins(1,4,5)P_3_ / Ins(3,5,6)P_3_ is reported as Ins(1,4,5)P_3_. In the InsP_3_ pool, four major isomers of InsP_3_, i.e Ins(1,2,3)P_3_, Ins(1,2,6)P_3_, Ins(1,4,5)P_3_ and Ins(2,4,5)P_3_ (**Fig. S1B**). Additionally, multiple isomers were detected for InsP_4_ (**Fig. S1C**) and InsP_5_ (**Fig. S1D,E**). Among the most abundant InsP_4_ isomers were Ins(1,4,5,6)P_4_ and Ins(1,2,4,6)P_4_/Ins(2,3,4,6)P_4_. For InsP_5_, Ins(1,2,4,5,6)P_5_ was the most abundant isomer, consistent with speciation in *Arabidopsis* (Riemer et al., 2021).

**Figure 3.**
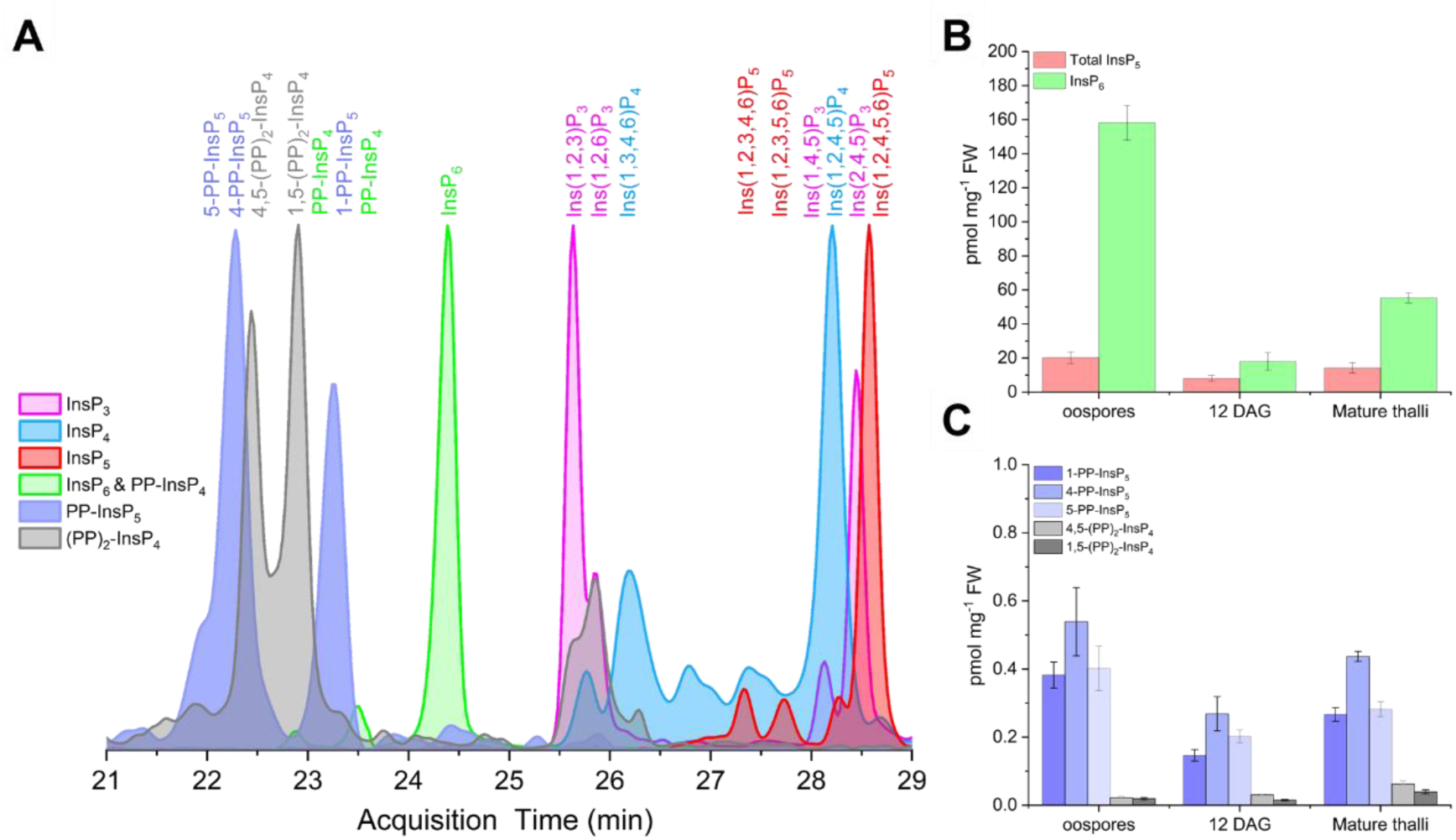
Detection and developmental profiling of inositol phosphates and pyrophosphates in *Chara braunii*. **A)** Detection and separation of PP-InsPs and InsPs from *Chara braunii* samples by CE-ESI-MS. Representative overlaid electropherograms showing resolution of pyrophosphorylated species and lower-order inositol phosphates. Background electrolyte (BGE): 35 mM ammonium acetate titrated with ammonium hydroxide to pH 9.7; CE voltage: 30 kV; CE current: 23 µA; injection: 10 nL. Peak identities were assigned based on migration time and co-migration with isotope-labeled references. Percent values indicate the relative distribution of positional isomers within each InsP class (e.g., InsP_3_, InsP_4_, PP-InsP_5_) and are normalized independently for each group; therefore, values are not comparable between different InsP classes. **B)** Quantification of total InsP_5_ and InsP_6_ concentrations across developmental stages of *Chara braunii.* InsP_6_ represents the dominant species at all stages, with highest abundance in spores and reduced levels during vegetative growth, whereas all three isomers of InsP_5_, i.e. Ins(1,2,4,5,6)P_5_, Ins(1,2,3,5,6)P_5_ and Ins(1,2,3,4,6)P_5_ remained detectable across development. **C)** Quantification of PP-InsPs across developmental stages. Relative levels of InsP_7_ isomers (5-PP-InsP_5_, 4-PP-InsP_5_ and 1-PP-InsP_5_) together with InsP_8_ are shown. For enantiomers, only a single possibility is indicated in all panels. All data represent mean ± SD; n = 3 biological replicates.

Quantitative profiling revealed pronounced stage-dependent differences across the InsP network. Phytic acid (InsP_6_) is the most abundant metabolite across all stages and shows substantial stage-dependent variation (**Fig. 3B**). Oospores, for example, had the highest InsP_6_ levels (158 pmol mg^-1^ fresh weight (FW)), which decreased almost 10-fold at 12 DAG (18 pmol mg^-1^ FW). In addition, low-abundance PP-InsP_4_ isomers, i.e. inositol pyrophosphates with only 6 phosphate groups, were detected during DAG stages, whereas these species were nearly absent in both oospores and mature thalli (**Fig. S1F)**.

We also analyzed PP-InsPs across development. Multiple PP-InsP_5_ isomers, including 1-PP-InsP_5_, 4-PP-InsP_5_, 5-PP-InsP_5_ and an unknown isomer (**Fig. S2G)**, were detected. 4-PP-InsP_5_ is the most abundant PP-InsP_5_ species across development (**Fig. 3C, Fig. S2H**), while 1-PP-InsP_5_ and 5-PP-InsP_5_ were detected in similar abundance (**Fig. 3C**). In addition, 1,5-(PP)_2_-InsP_4_ and 4,5-(PP)_2_-InsP_4_ were detected and assigned with stable isotope labeled references (Harmel et al., 2019; Desfougères et al., 2022). We noted the presence of at least another yet unassigned InsP_8_ isomer (grey peak at ca. 26 minutes in **Fig. 3A**). The 4,5-PP_2_-InsP_4_ isomer increased progressively during development, from 0.022 pmol mg^-1^ FW in oospores to 0.030 pmol mg^-1^ FW at 12 DAG and 0.062 pmol mg^-1^ FW in mature thalli (**Fig. 3C**).

Together, these data demonstrate that developmental changes in the inositol phosphate network of *Chara braunii* are substantial and involve a broad range of molecular species. In addition, the detection of a diverse array of isomers suggests the involvement of a complex and specialized enzymatic machinery. To investigate the molecular basis underlying this (developmental) regulation, we subsequently conducted a bioinformatic analysis to identify candidate kinase and phosphatase enzymes required in the synthesis, interconversion, and turnover of inositol phosphates in *Chara braunii*.

### Putative inositol polyphosphate metabolism-related genes in Chara braunii

Following the confirmation that various InsP and PP-InsP species were indeed produced in *Chara braunii*, we aimed to identify genes encoding potential enzymes involved in the InsP metabolism. An overview on putative candidates is given in **Table 1**.

**Table 1.** *Chara braunii* homologs of *A. thaliana* proteins involved in InsP and PP-InsP production. Putative proteins with sequence and structural similarity to selected, known plant enzymes involved in InsP metabolism are listed. The accession number of the respective query protein in *A. thaliana* is given, followed by the enzyme name, respective protein domains, the accession number of the best-matching protein in *C. braunii*, followed by the percentage of identical amino acids and the length of the sequence overlap (including gaps). Putative InsP enzymes from *C. braunii* were identified by sequence similarity using BLASTP (Altschul et al., 1990), or by similarities in domain architecture using INTERPRO (Blum et al., 2025). Note that neither the biochemical activities, nor the sequence identities *in vivo* have been experimentally assessed for the listed *Chara* proteins. Shown results were restricted to top three BLASTP results (or less if applicable), with a complete list available in **Supplementary Table S1**. Asterisks indicate that regulation was observed in transcriptomic analyses, detailed in the text.

| Accession no. | Enzyme | Domain | Chara homolog | Sequence identity | Overlap |
| --- | --- | --- | --- | --- | --- |
| **Kinases mediating the synthesis of InsPs** |  |  |  |  |  |
| NP_196354 | IPK2a | IPK_sf | g44488* | 39% | 286/205 |
|  |  | IPK_sf | g16917 | 34% | 286/514 |
| NP_001331991 | IPK2b | IPK_sf | g44488* | 39% | 300/205 |
|  |  | IPK_sf | g16917 | 36% | 300/514 |
| Q93YN9 | IPK1 | IP5_2-K_N_lobe | g22893 | 33% | 451/1350 |
| Q9SBA5 | ITPK1 | ITPK1_N | g23313* | 44% | 319/340 |
|  |  | InsP(3)kin | g54337* | 32% | 319/325 |
|  |  | Ins134_P3_kin | g3607 | 27% | 319/638 |
| O81893 | ITPK2 | ITPK1_N | g23313* | 56% | 391/340 |
|  |  | InsP(3)kin | g54337* | 36% | 391/325 |
|  |  | Ins134_P3_kin | g3607 | 25% | 391/638 |
| F4J8C6 | VIH1 | His_PPase superfam | g24415* | 68% | 1049/1524 |
| Q84WW3 | VIH2 | His_PPase superfam | g24415* | 68% | 1050/1524 |
| **Phosphoinositide-specific phospholipases** |  |  |  |  |  |
| Q39033 | PLC2 | PI-PLC Y-box | g36522* | 44% | 581/1860 |
| Q9LY51 | PLC7 | PI-PLC Y-box | g36522* | 43% | 584/1860 |
| Q944C1 | PLC4 | PI-PLC Y-box | g36522 | 43% | 597/1860 |
| **Phosphohydrolases** |  |  |  |  |  |
| Q9ZVN4 | DSP1 | Prot-tyrosine_phosphatase-like | g23885 | 69% | 146/215 |
|  |  |  | g29374* | 41% | 406/215 |
|  |  |  | g30804* | 32% | 235/215 |
| Q84MD6 | DSP2 | Prot-tyrosine_phosphatase-like | g23885 | 54% | 146/257 |
|  |  |  | g29374* | 33% | 406/257 |
|  |  |  | g19745* | 39% | 294/257 |
| Q681Z2 | DSP3 | Prot-tyrosine_phosphatase-like | g23885 | 51% | 146/203 |
|  |  |  | g29374* | 39% | 406/203 |
|  |  |  | g30804* | 36% | 235/203 |
| Q940L5 | DSP4 | Prot-tyrosine_phosphatase-like | g23885 | 66% | 146/198 |
|  |  |  | g29374* | 41% | 406/198 |
|  |  |  | g19745* | 43% | 294/198 |
| Q9FFD7 | DSP5 | Prot-tyrosine_phosphatase-like | g23885 | 57% | 146/204 |
|  |  |  | g29374* | 36% | 406/204 |
|  |  |  | g30804* | 36% | 235/204 |
| Q9LE73, Q9ZU95, | NUDT4/17 /18/21 | NUDIX_hydrolase_CS | g45972* | 34-37% | 566/176-198 |
| Q9LQU5,<br>Q8VY81 | (subclade<br>I) |  |  |  |  |
| Q93ZY7,<br>Q52K88,<br>Q9LHK1 | NUDT12/1<br>3/16<br>(subclade<br>II) | NUDIX_hydrolase<br>_CS | g45972* | 35-42% | 566/180-203 |

The first class of relevant enzymes are kinases mediating the synthesis of InsPs. There are four distinct inositol phosphate kinase families ubiquitous across eukaryotes: (i) inositol polyphosphate kinase IPK (which can be divided into subgroups IP3-3K, IPMK/Ipk2 and IP6K, the latter missing throughout the green lineage), (ii) inositol pentakisphosphate 2-kinase IPPK/IPK1, (iii) inositol 1,3,4-trisphosphate 5/6-kinase ITPK1, as well as (iv) diphosphoinositol pentakisphosphate kinase PPIP5K/Vip1/VIH (Miller et al., 2005; González et al., 2010; Shears and Wang, 2019; Randall et al., 2020; Laha et al., 2021b).

The second class of considered enzymes are phosphoinositide-specific phospholipases C (PI-PLCs), which catalyze the hydrolysis of phosphatidylinositol 4,5-bisphosphate into the second messengers inositol 1,4,5-trisphosphate (IP_3_) and 1,2-diacylglycerol. PI-PLCs are ubiquitous in eukaryotes, the *Arabidopsis* genome contains nine *AtPLC* genes. These AtPLCs were shown to be involved in responses to environmental stimuli, expressed in distinct tissues and evolved through multiple gene duplication events (Otterhag et al., 2001; Tasma et al., 2008).

The third class of relevant enzymes are phosphohydrolases that convert PP-InsP into their InsP precursors. These are members of the plant and fungal atypical dual-specificity phosphatase family (PFA-DSP), and certain NUcleoside DIphosphate linked to some other moiety X (NUDIX) hydrolases. PFA-DSPs are known for their broad substrate specificity as a mechanistically diverse superfamily (Glasner et al., 2006; Gunawardana et al., 2009). *Arabidopsis* possesses five PFA-DSP homologs, targeting 5-β-P of 5-InsP_7_ and 1,5-InsP_8_ (Gaugler et al., 2022; Ghosh et al., 2025). A family of NUDIX-type hydrolases (NUDTs) was characterized in *Arabidopsis* to catalyze the dephosphorylation of PP-InsP_5_ and (PP)_2_-InsP_4_ variants (Laurent et al., 2024; Ghosh et al., 2025; Schneider et al., 2025).

### Enzyme candidates for the synthesis of InsPs

In *Arabidopsis*, the IPK2 proteins phosphorylate 1,4,5-InsP_3_ sequentially at the 6-OH and 3-OH positions to generate 1,3,4,5,6-InsP_5_. The last step of InsP_6_ synthesis is then catalyzed by IPK2 (**Fig. 1**). Recent work showed that both AtIPK2α and AtIPK2β isoforms also govern the synthesis of 4-PP-InsP_5_ via phosphorylation of InsP_6_ and are critical in plant acclimation to heat stress. This mode of InsP_7_ synthesis, as well as heat stress response mediation, are conserved within the bryophyte *Marchantia polymorpha* (Cridland, 2022; Yadav et al., 2025).

*M. polymorpha* possesses a single enzyme, called MpIPMK (Yadav et al., 2025). Two putative IPMK homologs, g16917 and g44488, were identified in *Chara* at 34% to 39% sequence identity with MpIPMK. Structural comparison supported the relatedness of g16917 (ipTM score of 0.57, pTM score of 0.61; **Fig. 4A**) and of g44488 to MpIPMK (ipTM score of 0.47, pTM score of 0.6; **Fig. 4B**).

**Figure 4.**
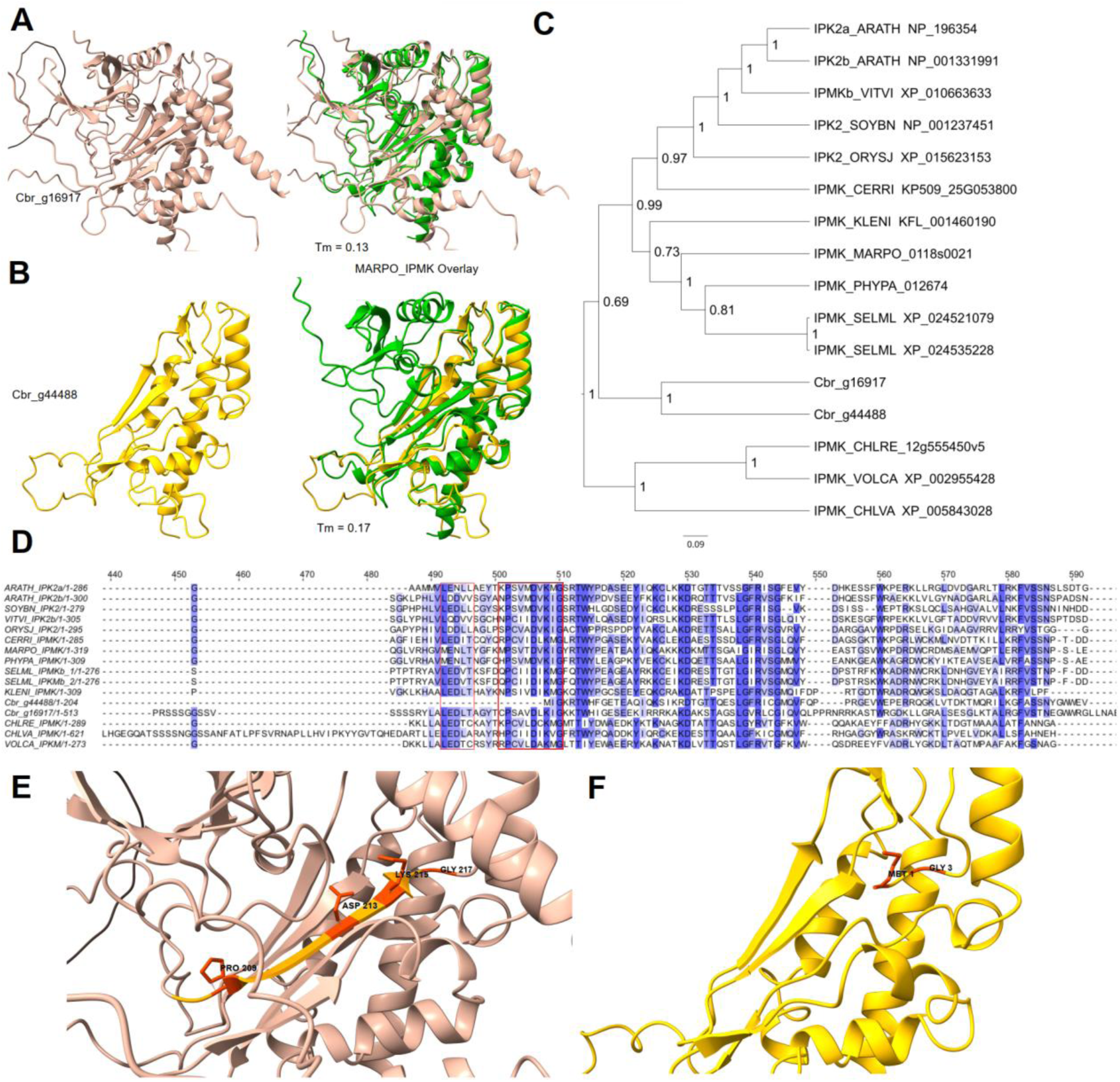
Domain structure, sequence comparison and phylogenetic relationships of *Chara braunii* IPMK candidates. **A)** Structural model overview of IPKM candidate g16917 (ipTM = 0.57, pTM = 0.61) and overlay with MpIPMK (Tm = 0.1291). **B)** Structural model overview of IPKM candidate g44488 (ipTM = 0.47, pTM = 0.6) and overlay with MpIPMK (Tm = 0.1720). Models for *Marchantia* IPMK and *Chara* were obtained via UniProtKB and visualized via ChimeraX. **C)** Bayesian inference of *Chara braunii* IPMK homologs. **D)** Multiple alignment of the IPMK central sequences from AtIPK2a/b homologs in selected viridiplantae model organisms. The alignment was generated by M-coffee and visualized using JalView v.2.11.2.7 (default colored percentage identity). Conserved LxxLL protein recognition and PxxxDxKxG catalytic motifs are boxed in red. The complete alignment is shown in Supplemental **Fig. S1. E)** Detailed view of conserved catalytic active site of g16917 and **F)** g44488. Motif and conserved residues are highlighted. Dataset of sequences for alignment and phylogenetic tree was based on Yadav et al. (2025).

Their phylogenetic analysis indicated a shared evolutionary lineage across the Viridiplantae, with both *Chara braunii* homologs at an ancestral position to all compared land plant homologs, while the homologs from non-streptophyte green algae *Chlorella*, *Chlamydomonas* and *Volvox* formed a separate clade (**Fig. 4C**). This result is consistent with the previously reported evolution of IPK2-type proteins across the viridiplantae from a single joint ancestry (Yadav et al., 2025). A multiple sequence alignment (**Fig. 4D**) showed the partial conservation of the protein recognition motif LxxLL (Plevin et al., 2005) in g16917 and of the PxxxDxKxG catalytic motif found within the InsP binding domain across all eukaryote branches (**Fig. 4E**). The IPMK homolog g44488, however, lacks an N-terminal protein recognition motif, and part of the conserved catalytic motif, K-to-M1 (lysine to methionine), but retaining the conserved glycine residue G3 (**Fig. 4F**). These findings suggest a putative loss or shift of function for g44488.

Furthermore, we identified one *Chara* IPK1 homolog, g22893 (GBG76676), at 33% sequence identity and more than twice the length of the *Arabidopsis* query protein (NP_568613). Domain predictions identified both a ribonuclease H superfamily-like (IPR012337) and Tf2-1-like SH3-like domain (IPR056924) within the N-terminal half of the protein, followed by an inositol-pentakisphosphate 2-kinase family associated region (IPR009286). Combined with an identified N-terminal RT_LTR domain, this suggests an instance of retrotransposon insertion.

Lacking IP6K/Kcs1-type proteins to synthesize PP-InsPs, *A. thaliana* utilizes its two Vip1 orthologs VIH1/VIP2L and VIH2/VIP1L (Laha et al., 2015) to phosphorylate InsP_6_ to 1-InsP_7_ and 5-InsP_7_ to 1,5-InsP8, as well as to hydrolyze 1-InsP_7_ and 5-InsP_7_ back to InsP_6_ (Riemer et al., 2022). Our query identified a single VIH homolog in *Chara*, g24415 (GBG59071), with a sequence identity of 68% to both the *A. thaliana* VIH1 and VIH2. Phylogenetic analysis clustered the sequence of the single *C. braunii* VIH homolog ancestral to all other compared homologs from the viridiplantae, with distinct clustering of the two *Arabidopsis* homologs indicating their origin from a likely recent gene duplication. Yeast Vip1, as an outgroup, still shares a sequence identity of 35% with *Arabidopsis* VIP1L (**Fig. 5A**). These kinases usually possess a dual-domain structure with an N-terminal ATP-grasp kinase domain, as well as a C-terminal histidine phosphatase superfamily-like domain (Laha et al., 2015; Zhu et al., 2019; Laha et al., 2021b; Schneider et al., 2025). The N-terminal kinase domain is conserved in *M. polymorpha* (Laurent et al., 2024; Ghosh et al., 2025; Rana et al., 2025). The diphosphoinositol pentakisphosphate kinase 2 N-terminal domain (IPR040557), belonging to the ATP-grasp superfamily (Desai et al., 2014; Raia et al., 2025), appears partially conserved within the *Chara* homolog, particularly the aspartic acid residue (D246) required for kinase activity has been retained alongside the arginine residue (R339) required for ligand binding (Williams et al., 2015) (**Fig. 5**, complete multiple sequence alignment in Supplemental **Fig. S2**). The presence of only one VIH-homolog in *Chara* is interesting, as we have observed at least three distinct InsP_8_ isomers (**Fig. 3**), suggesting that this homolog has either a broad substrate promiscuity or that other less-closely related enzymes are also producing InsP_8_ isomers.

**Figure 5.**
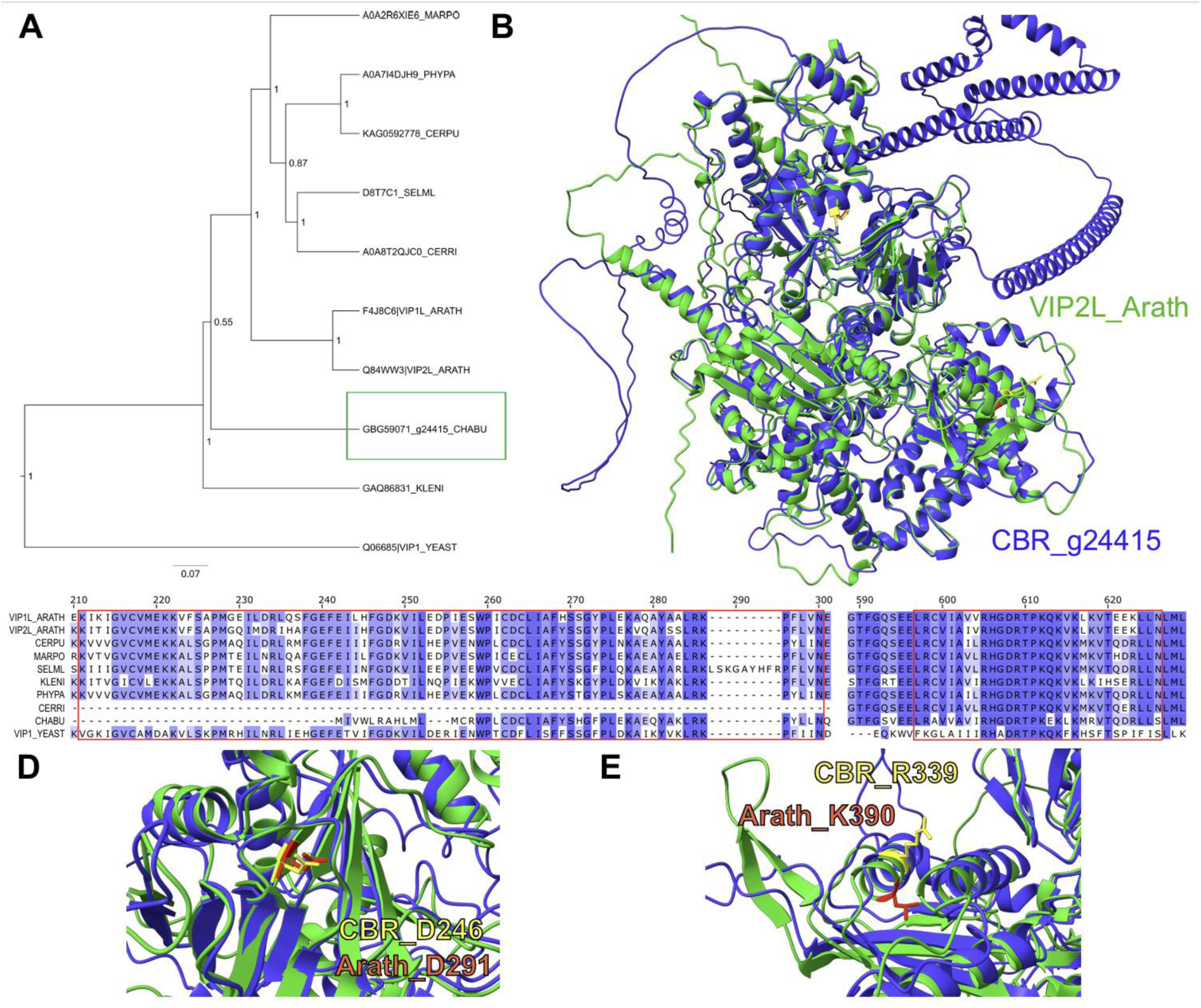
Phylogenetic relationships, domain structure and alignment of VIP candidates in *Chara braunii.* **A)** Bayesian inference of the phylogenetic relatedness of a *Chara braunii* VIH candidate (boxed in green) and homologs from land plants, Klebsormidium nitens (GAQ86831_KLENI) and yeast. **B)** Structural model of VIH candidate g24415 (pTM = 0.59) and overlay with AtVIP2L (Tm = 0.1514). Models for *Arabidopsis* VIP2L and *Chara* were obtained via UniProtKB and visualized via ChimeraX. **C)** Alignment of M-coffee aligned amino acid sequences was generated using JalView v.2.11.2.7 (default colored percentage identity). Conserved diphosphoinositol pentakisphosphate kinase 2 N-terminal domain and histidine acid phosphatases phosphohistidine signature are highlighted in red boxes. **D)** Detailed view of conserved asparagine residue (D291 in *Arabidopsis*) required for kinase activity in PPIP5K PH domain and **E)** arginine residue required for ligand binding (lysine 390 in *Arabidopsis*) of g24415. Conserved residues are each highlighted.

*Arabidopsis* ITPK1/2 kinases, belonging to the ATP-grasp fold family of proteins, further generate 5-InsP_7_ by phosphorylating InsP_6_, while their homologs ITPK3/4 notably do not (Laha et al., 2019; Riemer et al., 2022). We identified three putative ITPK homologs in *Chara*, g23313 (GBG76982), g54337 (GBG92082) and g3607 (GBG68908). Interestingly, g23313 shares the highest sequence identity with both queries, 44% for AtITPK1 and 56% for AtITPK2 (**Table 1**), but also with AtITPK3 with 62% sequence identity. Phylogenetic analysis indicated a subdivision of *Chara* ITPKs into distinct groups: Candidate g3607 clustered with *Arabidopsis* ITPK4, rice ITPK6 and *Marchantia* ITPK4, while g23313 clustered with *Arabidopsis* and *Marchantia* ITPK2/3. This finding is in line with previous studies (Laha et al., 2019; Pullagurla et al., 2025) indicating the subdivision of ITPKs into evolutionary conserved, distinct groups. Our results show that this subdivision of ITPK1-type kinases across the Viridiplantae includes the *Chara* homologs.

The third homolog, g54337, meanwhile appeared distinct from the ITPK4 cluster, yet lower branching in relation to the ITPK2/3 and ITPK1 clusters (**Fig. S3**, for the complete multiple sequence alignment, see **Fig. S4**).

### A single phosphoinositide-specific phospholipase candidate

We identified a single potential *Chara* PLC homolog, g36522 (GBG82993). The protein G36522 exhibited 44%, 43% and 43% sequence identity with the AtPLC2, AtPLC7 and AtPLC4 homologs, while the identity to the other 6 enzymes in *Arabidopsis* was lower. This PLC homolog contains a conserved N-terminal PLC-like phosphodiesterase (IPR001192) and a C-terminal C2-superfamily-like domain (IPR035892). However, the putative protein is roughly thrice the size of queried *Arabidopsis* proteins, caused by an additional C-terminal ribonuclease H superfamily-like (IPR036397), a DNA/RNA polymerase superfamily-like (IPR043502), as well as an RT_LTR domain in between the aforementioned PLC-like and C2-like domains, suggesting a retrotransposon insertion event. The presence of single PLC suggests that PLC-mediated InsP_3_ production may be differentially regulated or limited compared to algae and land plants.

### Assignment of PP-InsP-converting phosphohydrolases

To assure homeostasis of hormone signaling, nutrient sensing and plant immunity, phosphohydrolases are needed to convert PP-InsPs back into their InsP precursors. *Arabidopsis* possesses five members of the PFA-DSP family (AtDSPs), targeting 5-InsP_7_ and 1,5-InsP_8_ by cleaving their 5-β phosphate (Gaugler et al., 2022; Ghosh et al., 2025).

In *Chara* we identified the same number, i.e. five putative DSP homologs to these *Arabidopsis* enzymes (g23883, g23885, g29374, g30804 and g19745). From these, g23885 (GBG77436) shares the highest sequence identities with all 5 AtDSPs, between 69% and 51%, respectively.

The *A. thaliana* genome also contains 28 NUDIX family genes (Kraszewska, 2008; Yoshimura and Shigeoka, 2015), which have been classified into 11 individual families with distinct, assigned functions (Gunawardana et al., 2009). Two distinct subclades of NUDIX-type hydrolases (NUDTs) with distinct and specific PP-InsP pyrophosphatase activities were recently identified: Subclade I enzymes (AtNUDT4/17/18/21) mainly target 4-InsP_7_, while subclade II (AtNUDT12/13/16) enzymes preferentially target 3-InsP_7_ (Laurent et al., 2024; Schneider et al., 2025). In *Chara*, we identified 15 putative homologs to the AtNUDT genes in *A. thaliana.* Four of these identified candidates, g49120, g49122, g49131 and g49132 share putative retroelements. Only one of these homologs, the *C. braunii* NUDT gene g45972 was associated by BLASTP with the two diphosphoinositol polyphosphate phosphohydrolase subclades previously identified in *A. thaliana*, sharing between 42% and 34% sequence identity, respectively. Phylogenetic analysis revealed strong clustering support for the NUDT encoded by g45972 with subclade II (**Fig. 6**).

**Figure 6.**
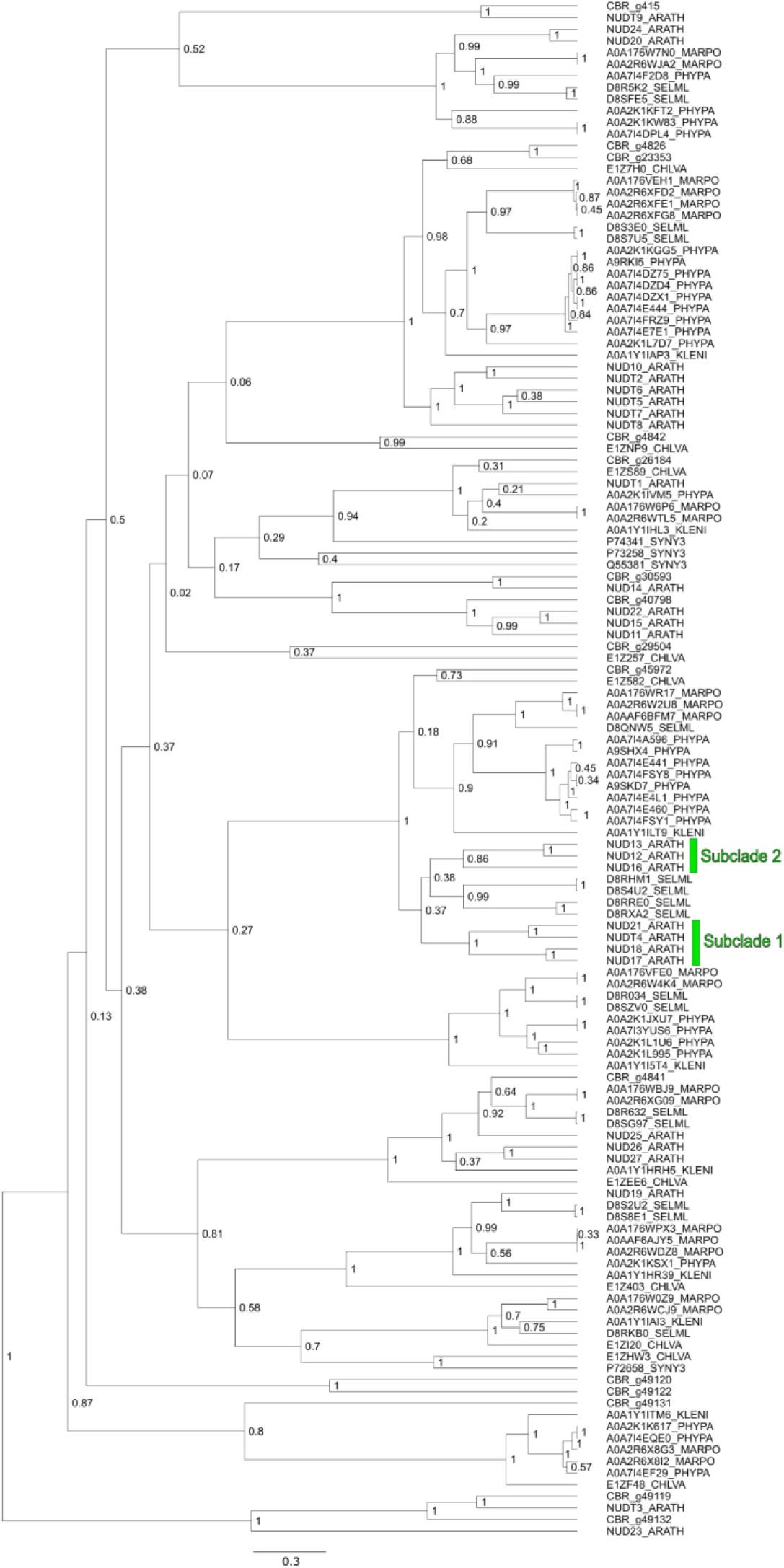
Phylogenetic relationships of *Chara braunii* Nudix candidates. Bayesian inference of *Chara braunii* NUDT homologs (in comparison to *A. thaliana*. Dataset of sequences for alignment and phylogenetic tree generation was based on (Schneider et al., 2025).)

Together, these data suggested that the inositol phosphate metabolism in *Chara braunii* is linked to signalling, and stress-related pathways. We therefore next examined whether environmental conditions influence the InsP network by analysing metabolite levels under stress conditions.

### Stress-dependent changes in InsP metabolism in Chara braunii

Analysis of previously performed RNA-seq experiments (Heß et al., 2023b; Heß et al., 2023a; Heise et al., 2025) indicated that several of the here assigned enzymes were differentially expressed during different environmental conditions (**Fig. S5**). To investigate whether environmental conditions influence InsP metabolism, we next analyzed InsP and PP-InsP levels after heat stress, prolonged light and dark treatments and after transfer to deionized water to exclude the effects of possible InsP molecules leaked from the soil substrate (**Fig. 7 and Fig. S7 and S8**). The cultures were sampled at 4, 8, 12, and 16 DAG (**Fig. 2B**) for CE-MS analysis. Thalli at 16 DAG were subsequently subjected to prolonged light or darkness (**Fig. 2B**). Mature thalli grown under 16/8 h light/dark standard conditions were exposed to defined stress treatments to assess condition-dependent changes in the InsP network. First, we analyzed the effect of the growth condition by comparing the thalli grown in deionized water for 3 days with developing thalli at 4 DAG (**Fig. 2C**). Thalli grown in deionized water showed a reduction in total InsP_5_ levels, which decreased approximately twofold (7.8 to 3.9 pmol mg^-1^ FW), whereas InsP_6_ showed only modest variation. A pronounced reduction was observed for PP-InsP_5_ species, with 1-PP-InsP_5_, 4-PP-InsP_5_, and 5-PP-InsP_5_ all ca. 10-fold decreased. Overall, transfer to deionized water lowered InsP_5_ and PP-InsP_5_ levels, while InsP_6_ concentrations remained comparatively stable (**Fig. S7A, B)**.

**Figure 7.**
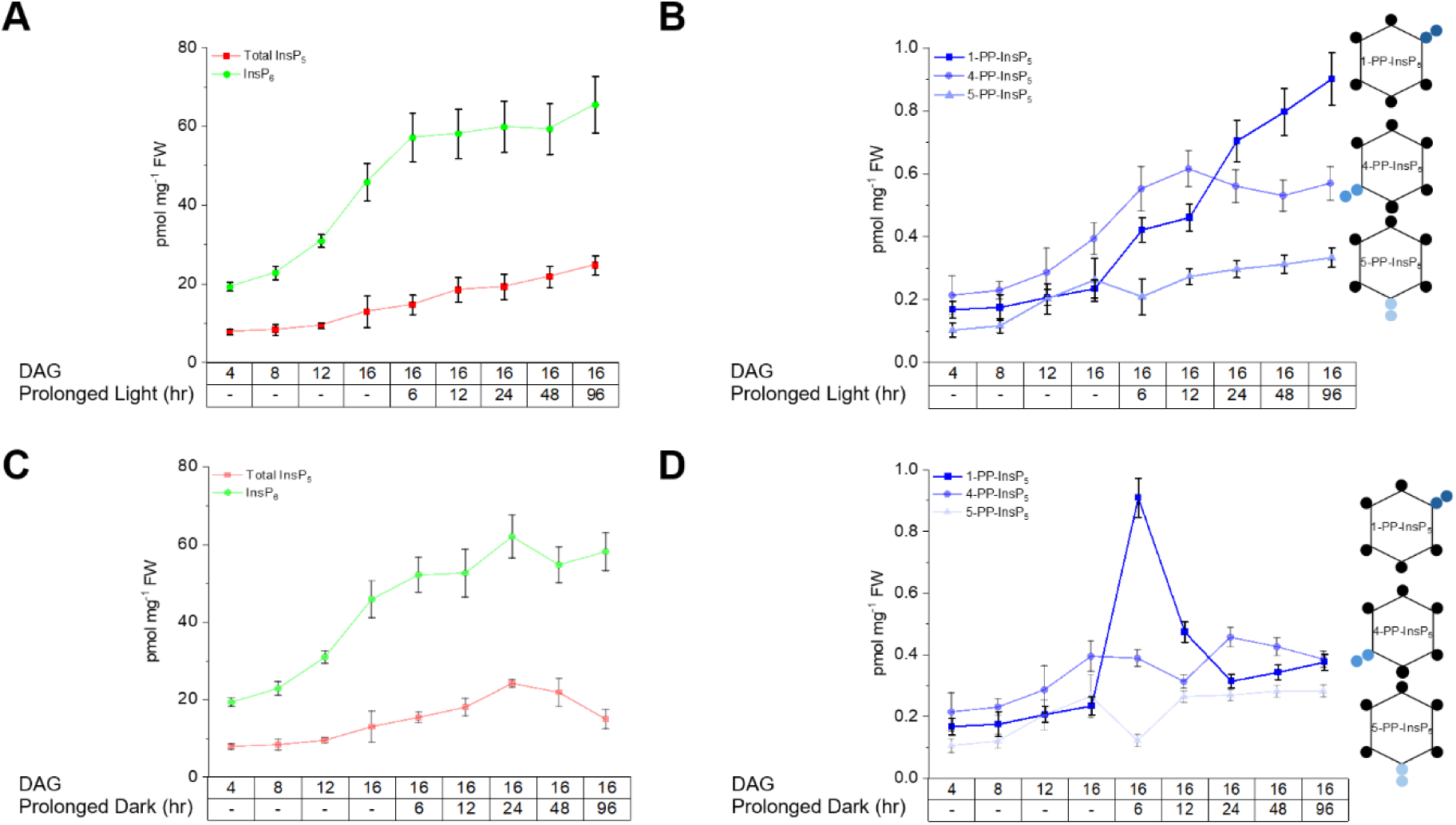
Light- and dark-dependent remodeling of inositol phosphate metabolism in *Chara braunii*. **A)** Quantification of InsP_5_ and InsP_6_ levels during prolonged growth under 16/8 h light/dark. **B)** Quantification of PP-InsP_5_ isomers during prolonged growth under 16/8 h light/dark. Illumination was provided by lateral white light (30 – 70 μmol photons m ^-2^ s ^-1^). **C)** Quantification of InsP_5_ and InsP_6_ levels during prolonged dark treatment. **D)** Quantification of PP-InsP_5_ isomers during prolonged dark exposure. Data represent mean ± SD; n = 3 biological replicates.

Further we examined the effect of 48 h exposure to 32^°^C (heat stress) on the 16 DAG *Chara* samples (**Fig. 2D**). Pronounced changes were observed after 12 h of heat exposure (**Fig. S7C, D**). Relative to 16 DAG controls, heat-stressed samples showed a ca. 1.3-fold increase in total InsP_5_ and InsP_6_ levels (**Fig. S7C, D**). Inositol pyrophosphate species showed differential responses, with a ca. 1.3-fold modest increase in 1-PP-InsP_5_. In contrast, 4-PP-InsP_5_ shows a marked decrease (**Fig. S7C, D**). Together, these results indicate that short-term heat stress promoted a pronounced reduction in 4-PP-InsP_5_, in line with previous results obtained in *Marchantia* (Yadav et al., 2025).

We next analyzed the effect of continued growth under 16/8 h light/dark after 16 DAG. This treatment resulted in the progressive accumulation of some InsPs. InsP_6_ increased from 45.8 pmol mg^-1^ FW at 16 DAG to 57.1 pmol mg^-1^ FW after 6 h and reached 65.5 pmol mg^-1^ FW after 96 h of continued growth at light/dark conditions. Similarly, total InsP_5_ increased from 12.9 pmol mg^-1^ FW to 24.7 pmol mg^-1^ FW over the same time-period **(Fig. 7A, S8A)**. Inositol pyrophosphates showed more pronounced light-dependent responses. In particular, 1-PP-InsP_5_ increased ca. 4-fold after 96 h of prolonged light compared to 16 DAG, becoming the dominant PP-InsP **(Fig. 7B).**

In contrast to light exposure, dark treatment produced dynamic and transient remodeling of InsP species. Relative to 16 DAG, InsP_6_ increased ca. 1.4-fold at 24 h, followed by moderate variation at later time points **(Fig. 7C)**. Total InsP_5_ showed a similar transient increase of ca. 2-fold after 24 h, before declining at prolonged dark exposure **(Fig. 7C, S8B)**. In contrast, the PP-InsPs - specifically again 1-PP-InsP_5_ - showed more pronounced transient responses. 1-PP-InsP_5_ spiked by ca. 4-fold after 6 h, followed by a rapid decline up to 12 h (0.474 pmol mg^-1^ FW) returning to baseline levels during the following 24-96 h **(Fig. 7D)**. Together, these results indicate that in particular 1-PP-InsP_5_ was most responsive to the prolonged light and dark incubations. This provides a rare example where specifically this PP-InsP isomer alone is mostly responsive to an external stimulus.

## Discussion

The transition from water to land challenged the physiology of early land plants in multiple ways, by desiccation, enhanced exposure to UV, different light regimes and generally, more rapidly changing environmental conditions. Recent findings have now started to illuminate the critical role PP-InsPs might have played in these processes as second messengers. PP-InsPs are involved in P_i_ homeostasis, participate in phytohormone perception and crosstalk, and contribute to (a)biotic stress responses and acclimation (reviewed by (Ghosh et al., 2025; Giehl and Schaaf, 2026)).

In this study, we aimed to identify and bioinformatically characterize homologs to known plant proteins involved in InsP metabolism in *Chara braunii*. Our metabolic profiling demonstrates that *Chara braunii* has a developmentally responsive InsP network that extends from lower InsPs to inositol pyrophosphate species (**Fig. 2A**). Although the detection and resolution of PP-Insp isomers continues to be a challenge due to their low abundance and transient nature (Laha et al., 2021a; Qiu et al., 2023), we provide data that hints at a degree of metabolic complexity that involves currently unknown interconversion routes potentially requiring enzymes that remain to be identified. For example, we detected at least four InsP_7_ and three InsP_8_ species, while only a single VIH homolog is identified in the genome (**Fig. 5**).

As a representative of algal progenitors to embryophytes, the *Chara* model is now situated among a variety of species across the green lineage for which genes encoding InsP kinases and phosphatases, as well as a diverse array of produced InsP species, were identified (Laha et al., 2021b; Chen et al., 2023; Morales-Pineda et al., 2023; Xiong et al., 2024; Bedera-García et al., 2025; Pullagurla et al., 2025; Yadav et al., 2025). In light of the growing literature, it has been hypothesized that the expansion and diversification of InsP metabolic networks may have contributed towards the green adaption to terrestrial environments (Ghosh et al., 2025).

Recent efforts unveiled the conservation of crucial InsP biosynthesis enzymes between bryophytes and embryophytes, like the conserved regulation of P_i_ homeostasis by ITPK1-derived PP-InsPs (Pullagurla et al., 2025), the InsP_8_ synthesis and developmental mediation of VIH (Rana et al., 2025), and the PP-InsP synthase activity of IPK2 proteins (Yadav et al., 2025) in *Marchantia polymorpha*. Our data supports the evolutionary conservation of these crucial enzymes across the Viridiplantae. Cursory inspection of putative InsP regulatory networks within *Chara* additionally revealed connections to phytohormone signaling, transcriptional regulation at the chromatin level and energy metabolism. Future more comprehensive inference of co-expression gene regulatory networks across multiple RNA-seq datasets (Dadras, 2024) will undoubtedly help to unravel their breadth and evolutionary conservation across the green lineage.

Our data provides supporting evidence for the conservation of crucial InsP biosynthesis pathways, provides a foundation for further inquiries and supports the hypothesis that expansion and diversification of InsP metabolic networks may have contributed to green adaption towards terrestrial environments. Notably, we identify conditions (light/dark stress), where 1-PP-InsP_5_ becomes the dominant isomer and appears to be dynamically regulated. Future research is now needed to investigate the synthesis of InsPs *in vivo,* characterize the function of *Chara* enzyme candidates and examine their impact on the regulation of equivalent physiological processes in streptophyte algae.

## Material and Methods

### Chara cultivation and treatments

The streptophyte freshwater alga *Chara braunii* S276 was cultivated as previously described (Heß et al., 2023b; Holzhausen et al., 2024). Standard cultivation vessels were filled with a tertiary layer of quartz sand, lime and sieved compost under deionized water. All components were double-autoclaved. Algae were grown at 22°C under long-day conditions in a 16 h light/8 h dark cycle under lateral white light illumination (30 – 70 μmol photons m ^-2^ s ^-1^, Lumilux L36W/840, OSRAM). Mature oospores were collected from the sediment surface via pipette and stored in deionized water under darkness at 4 °C for at least six months before germination induction or InsP extraction (**Fig. 2A**).

To evaluate the shift of InsP levels under environmental stress in *Chara braunii*, whole thalli were grown from harvested oospores rinsed with double-autoclaved deionized water under standard conditions for 42 days. Liquid medium was exchanged once after two weeks. Thallus samples under standard conditions were periodically collected during growth, in triplicates, after 26 d, 30 d, 34 d and 38 d, with the last sampling timepoint serving as t0 for subsequent Light/Dark illumination stress experiment. Remaining culture vessels were divided into control (standard conditions) and dark (wrapped in aluminum foil) treatment. Algal samples were taken after 6 h, 12 h, 24 h, 48 h and 96 h after transfer to the respective conditions, frozen in liquid nitrogen and stored at -80°C. In parallel, liquid medium samples were collected, frozen at -20°C and stored for later processing (**Fig. 2B**). Additionally, vegetatively propagated, mature *Chara* thalli grown for at least 30 days were either transferred to deionized water for three days before collecting liquid and thallus samples (**Fig. 2C**), or kept at 32°C for three days (**Fig. 2D**). Both deionized water and heat stress experiments were performed under otherwise standard growth conditions. After harvesting, samples were frozen in liquid nitrogen and stored at -80°C until processed.

### Bioinformatic analyses and use of LLMs

A set of *Arabidopsis thaliana* proteins, involved in InsP and PP-InsPs metabolism, were selected based on their characterization and annotation, and their sequences collected from UniProt (The UniProt Consortium, 2025). Protein-protein BLAST searches (Altschul et al., 1990) were performed to identify potential homologous genes in *Chara braunii* by sequence similarity and coverage, paired with INTERPROScan queries (Blum et al., 2025) to identify similarities in domain architecture. For subsequent phylogenetic evaluations, the initial blastp query was extended to identify homologous genes within representative species across the Viridiplantae.

Tertiary structures of query proteins were predicted using Alphafold3 (Abramson et al., 2024) unless crystal- or NMR-resolved structures were available, and aligned using ChimeraX (Meng et al., 2023). To investigate topological similarity between protein structures, Tm scores were calculated via web application (https://aideepmed.com/TM-score/) (Zhang and Skolnick, 2004).

Multiple sequence alignments (MSA) were generated using M-coffee (Notredame et al., 2000; Di Tommaso et al., 2011) and analyzed using BEAST 2.7.8 (Bouckaert et al., 2019). Calculations were performed under standard BEAUTi settings utilizing the yule model tree prior, Blosum62 as substitution model and MCMC chain length of 1e7, with logged parameters every 1e4 steps. The resulting dendrograms were assessed with Tracer v1.7.2, and maximum clade credibility trees generated via TreeAnnotator using 50% burnIn and a posterior probability limit of 0.5 for median node heights (Rambaut et al., 2018). Resulting dendrograms were visualized using FigTree v1.4.4 (Rambaut, 2018) before further incorporation using Inkscape v1.4 (The Inkscape Developers, 2024).

ChatGPT (OpenAI, GPT-4o) was used for language refinement.

### InsPs extraction and TiO₂ enrichment

To extract InsPs, we used 100–200 mg (wet weight) of flash-frozen samples, making sure to keep everything ice-cold throughout the process. The samples were first homogenized in chilled 1 M perchloric acid (using about 800–1,000 μL per sample), followed by gentle rotation at 4°C to precipitate proteins and release metabolites. Then, samples were centrifuged at maximum speed (≥16,000 × g) for 15 minutes at 4 °C. The clear supernatant, which contains the soluble inositol phosphates, was carefully transferred to fresh tubes.

To enrich InsPs and pyrophosphates from these extracts, we used TiO₂ bead-based affinity purification (Wilson and Saiardi, 2018). Specifically, 5 mg of TiO₂ beads (Titansphere TiO_2_, 5 μm; GL Sciences) were first washed with water and then 1 M perchloric acid. These pre-washed TiO_2_ beads were added to the supernatant and rotated for 20 minutes at 4 °C. After binding, the beads were collected by centrifugation at 5000 × g and washed twice with ice-cold 1 M perchloric acid to remove any unspecifically binding contaminants.

To recover the InsPs, we incubated the beads twice with 10% ammonium hydroxide and rotated for 5 mins, pooled the eluates, and immediately dried them under vacuum using a SpeedVac concentrator. The dried samples were then stored at −20 °C, until reconstitution and analysis by CE-MS.

### Capillary Electrophoresis-Mass Spectrometry (CE-MS)

#### Chemicals and Reagents

For our experiments, we used high-quality chemicals to ensure accuracy. Methanol, ammonium acetate, 2-propanol, and a 10% ammonia solution were all sourced from Carl Roth. The ^13^C_6_-labeled inositol phosphates and pyrophosphates we used were kindly provided by Dorothea Fiedler at FMP Berlin (Harmel et al., 2019). ^18^O-labelled references were synthesized in house as described (Haas et al., 2022; Liu et al., 2025; Ritter et al., 2025).

#### Instrumentation and setup

All our analyses were done using an CE-ESI-QQQ system (Agilent 7100 CE with Agilent 6495C Triple Quadrupole and Agilent Jet Stream electrospray ionization source, adopting an Agilent CE-ESI-MS interface). For selected analyses, the CE system was alternatively coupled to a quadrupole time-of-flight (QToF) mass spectrometer using the same dedicated CE-MS adapter and sprayer kit. All measurements were performed in negative electrospray ionization mode. We used a bare fused-silica capillary (100 cm, 50 µm internal diameter). Before starting, we flushed the capillary with 1 M NaOH for 10 minutes and then with water for another 10 minutes.

Depending on what we were analysing, we used two different background electrolytes (BGEs).

BGE-1: 35 mM ammonium acetate, pH adjusted to 9.75 with ammonium hydroxide, for general profiling of inositol phosphates and pyrophosphates.

BGE-2: 40 mM ammonium acetate, pH adjusted to 9.08, for better separation of specific InsP_5_ and InsP_7_ isomers.

Before each run, we equilibrated the capillary with the chosen BGE for about 7 min (400 s). Samples (30 nL) were introduced by pressure injection (100 mbar for 15 seconds). For QToF measurements, we used a sheath liquid of water and 2-propanol (1:1, v/v) with continuous mass reference ions. For QqQ measurements, the sheath liquid was the same but without the reference ions. The sheath liquid flow was 10 µL per minute. We controlled the instruments and collected data using Agilent’s Mass Hunter Workstation software (version 10.1). Details about the instrument settings can be found in **Tables S2** and **S3**.

### Quantitative analysis

For quantitative analysis, dried TiO_2_-enriched extracts were reconstituted in deionized water prior to CE-MS measurement. Absolute and relative quantification of inositol phosphates was performed using stable isotope-labeled internal standards spiked directly into each sample before analysis.

InsP_5_, InsP_6_, PP-InsP_5_, and InsP_8_ were quantified using known amounts of their corresponding ^13^C_6_-labeled standards as internal references. For lower inositol phosphates (InsP_3_ and InsP_4_), for which isotopically labeled standards are not commercially available for all positional isomers, quantification was performed using spiked ^13^C_6_-InsP_6_ as internal standard.

When appropriate, pure ^3^C_6_-labeled Ins(1,2,3)P_3_ (Liu et al., 2023) was additionally used to validate InsP_3_ quantification and enable direct comparison across lower InsP species under identical CE–MS conditions. This strategy ensured consistent normalization across the inositol phosphate series despite differences in available isotopic standards.

Isotopic standards were spiked at concentrations specified in **Table S4**. These concentrations were selected to ensure accurate detection within the linear dynamic range of the instrument, enabling reliable and reproducible quantification.

## Acknowledgments

This work was supported by the Deutsche Forschungsgemeinschaft (DFG) through priority program 2237 “MAdLand”, http://madland.science, grant HE2544/18-2 to W.R.H., under Germany’s excellence strategy CIBSS, EXC-2189, Project ID 390939984, to H.J.J.; and a research grant JE 572/11-1 to H.J.J. A.S. and H.J.J. acknowledge funding from the Volkswagen Foundation (VW Momentum Grant 98604).

We thank Madeleine Tarika Maier and Pauline Mauser, Freiburg, for maintaining vegetative cultures of *Chara braunii* and providing technical support.

## Author Contributions

WRH and HJ conceptualized the study with input from DH and AS. AS performed and evaluated all chemical experiments. DH performed stress experiments and sampling with *Chara braunii* and carried out all bioinformatic analyses. WRH supervised the evaluation of the bioinformatics data. DH and WRH drafted the manuscript, and all authors were involved in writing the final manuscript.

## References

1. Abramson J, Adler J, Dunger J, Evans R, Green T, Pritzel A, Ronneberger O, Willmore L, Ballard AJ, Bambrick J, et al (2024) Accurate structure prediction of biomolecular interactions with AlphaFold 3. Nature 630: 493–500

2. Altschul SF, Gish W, Miller W, Myers EW, Lipman DJ (1990) Basic local alignment search tool. J Mol Biol 215: 403–410

3. Andrews M (1987) Phosphate uptake by the component parts of *Chara hispida*. British Phycological Journal 22: 49–53

4. Bedera-García R, García-Gómez ME, Personat JM, Couso I (2025) Inositol polyphosphates regulate resilient mechanisms in the green alga *Chlamydomonas reinhardtii* to adapt to extreme nutrient conditions. Physiologia Plantarum 177: e70089

5. Beilby MJ, Casanova MT (2014) The Physiology of Characean Cells. doi: 10.1007/978-3-642-40288-3

6. Bierenbroodspot MJ, Pröschold T, Fürst-Jansen JMR, de Vries S, Irisarri I, Darienko T, de Vries J (2024) Phylogeny and evolution of streptophyte algae. Annals of Botany 134: 385–400

7. Blum M, Andreeva A, Florentino LC, Chuguransky SR, Grego T, Hobbs E, Pinto BL, Orr A, Paysan-Lafosse T, Ponamareva I, et al (2025) InterPro: the protein sequence classification resource in 2025. Nucleic Acids Research 53: D444–D456

8. Bonnot C, Hetherington AJ, Champion C, Breuninger H, Kelly S, Dolan L (2019) Neofunctionalisation of basic helix−loop−helix proteins occurred when embryophytes colonised the land. New Phytologist 223: 993–1008

9. Bouckaert R, Vaughan TG, Barido-Sottani J, Duchêne S, Fourment M, Gavryushkina A, Heled J, Jones G, Kühnert D, Maio ND, et al (2019) BEAST 2.5: An advanced software platform for Bayesian evolutionary analysis. PLOS Computational Biology 15: e1006650

10. Braun M, Foissner I, Löhring H, Schubert H, Thiel G (2007) Characean Algae: Still a Valid Model System to Examine Fundamental Principles in Plants. In K Esser, U Löttge, W Beyschlag, J Murata, eds, Progress in Botany. Springer, Berlin, Heidelberg, pp 193–220

11. Chen Y, Han J, Wang X, Chen X, Li Y, Yuan C, Dong J, Yang Q, Wang P (2023) OsIPK2, a rice inositol polyphosphate kinase gene, is involved in phosphate homeostasis and root development. Plant Cell Physiol 64: 893–905

12. Cheng S, Xian W, Fu Y, Marin B, Keller J, Wu T, Sun W, Li X, Xu Y, Zhang Y, et al (2019) Genomes of subaerial Zygnematophyceae provide insights into land plant evolution. Cell 179: 1057–1067.e14

13. Corti B (1729-1813) (1774) Osservazioni microscopiche sulla tremella.

14. Couso I, Evans BS, Li J, Liu Y, Ma F, Diamond S, Allen DK, Umen JG (2016) Synergism between inositol polyphosphates and TOR kinase signaling in nutrient sensing, growth control, and lipid metabolism in *Chlamydomonas* . Plant Cell 28: 2026–2042

15. Couso I, Smythers AL, Ford MM, Umen JG, Crespo JL, Hicks LM (2021) Inositol polyphosphates and target of rapamycin kinase signalling govern photosystem II protein phosphorylation and photosynthetic function under light stress in Chlamydomonas. New Phytologist 232: 2011–2025

16. Cridland CA (2022) Elucidating the function of inositol pyrophosphate signaling pathways in *Arabidopsis thaliana*.

17. Dadras A (2024) Comparative analyses of stress-induced dynamics in global differential gene expression across more than 550 million years of streptophyte evolution. Georg-August-Universität Göttingen, Göttingen

18. Desai M, Rangarajan P, Donahue JL, Williams SP, Land ES, Mandal MK, Phillippy BQ, Perera IY, Raboy V, Gillaspy GE (2014) Two inositol hexakisphosphate kinases drive inositol pyrophosphate synthesis in plants. The Plant Journal 80: 642–653

19. Desfougères Y, Portela-Torres P, Qiu D, Livermore TM, Harmel RK, Borghi F, Jessen HJ, Fiedler D, Saiardi A (2022) The inositol pyrophosphate metabolism of *Dictyostelium discoideum* does not regulate inorganic polyphosphate (polyP) synthesis. Adv Biol Regul 83: 100835

20. Di Tommaso P, Moretti S, Xenarios I, Orobitg M, Montanyola A, Chang J-M, Taly J-F, Notredame C (2011) T-Coffee: a web server for the multiple sequence alignment of protein and RNA sequences using structural information and homology extension. Nucleic Acids Res 39: W13–17

21. Doege A, Becker R, Schubert H, van de Weyer K (2016) Bioindikation mit Characeen. Armleuchteralgen: Die Characeen Deutschlands. Springer, Berlin, Heidelberg, pp 97–137

22. Dong J, Ma G, Sui L, Wei M, Satheesh V, Zhang R, Ge S, Li J, Zhang T-E, Wittwer C, et al (2019) Inositol pyrophosphate InsP8 acts as an intracellular phosphate signal in *Arabidopsis*. Mol Plant 12: 1463–1473

23. Foissner I, Wasteneys GO (2014) Chapter Seven - Characean internodal cells as a model system for the study of cell organization. *In* KW Jeon, ed, International Review of Cell and Molecular Biology. Academic Press, pp 307–364

24. Fürst-Jansen JMR, de Vries S, de Vries J (2020) Evo-physio: on stress responses and the earliest land plants. Journal of Experimental Botany 71: 3254–3269

25. Gaugler P, Schneider R, Liu G, Qiu D, Weber J, Schmid J, Jork N, Häner M, Ritter K, Fernández-Rebollo N, et al (2022) *Arabidopsis* PFA-DSP-type phosphohydrolases target specific inositol pyrophosphate messengers. Biochemistry 61: 1213–1227

26. Ghosh R, Yadav R, Pullagurla NJ, Rana P, Laha D (2025) The expanding landscape of inositol phosphate signaling network in land plants. FEBS Letters. doi: 10.1002/1873-3468.70236

27. Giehl RFH, Schaaf G (2026) Metabolism, perception, andfunctions of inositol (pyro)phosphates in plants. Annu Rev Plant Biol. doi: 10.1146/annurev-arplant-070225-040214

28. Glasner ME, Gerlt JA, Babbitt PC (2006) Evolution of enzyme superfamilies. Current Opinion in Chemical Biology 10: 492–497

29. González B, Baños-Sanz JI, Villate M, Brearley CA, Sanz-Aparicio J (2010) Inositol 1,3,4,5,6-pentakisphosphate 2-kinase is a distant IPK member with a singular inositide binding site for axial 2-OH recognition. Proceedings of the National Academy of Sciences 107: 9608–9613

30. Gulabani H, Goswami K, Walia Y, Roy A, Noor JJ, Ingole KD, Kasera M, Laha D, Giehl RFH, Schaaf G, et al (2022) Arabidopsis inositol polyphosphate kinases IPK1 and ITPK1 modulate crosstalk between SA-dependent immunity and phosphate-starvation responses. Plant Cell Rep 41: 347–363

31. Gunawardana D, Likic VA, Gayler KR (2009) A comprehensive bioinformatics analysis of the nudix superfamily in *Arabidopsis thaliana*. International Journal of Genomics 2009: 820381

32. Haas TM, Mundinger S, Qiu D, Jork N, Ritter K, Dürr-Mayer T, Ripp A, Saiardi A, Schaaf G, Jessen HJ (2022) Stable isotope phosphate labelling of diverse metabolites is enabled by a family of 18 O-phosphoramidites. Angew Chem Int Ed Engl 61: e202112457

33. Harmel RK, Puschmann R, Nguyen Trung M, Saiardi A, Schmieder P, Fiedler D (2019) Harnessing 13C-labeled myo-inositol to interrogate inositol phosphate messengers by NMR. Chem Sci 10: 5267–5274

34. Heise CM, Heß DA, Walke P, Voß M, Schubert H, Hess WR, Hagemann M (2025) Evidence for a CO2-concentrating mechanism in the model streptophyte green alga *Chara braunii*. New Phytologist 247: 1218–1233

35. Heß D, Heise CM, Schubert H, Hess WR, Hagemann M (2023a) The impact of salt stress on the physiology and the transcriptome of the model streptophyte green alga *Chara braunii*. Physiologia Plantarum 175: e14123

36. Heß D, Holzhausen A, Hess WR (2023b) Insight into the nodal cells transcriptome of the streptophyte green alga *Chara braunii* S276. Physiologia Plantarum 175: e14025

37. Holzhausen A, Stingl N, Maier MT, Rensing SA, Schubert H (2024) Cultivation and germination induction of *Chara braunii* S276 : MadLand protocol collection. doi: 10.6094/UNIFR/245850

38. Irvine RF, Letcher AJ, Stephens LR, Musgrave A (1992) Inositol polyphosphate metabolism and inositol lipids in a green alga, Chlamydomonas eugametos. Biochem J 281: 261–266

39. Irvine RF, Schell MJ (2001) Back in the water: the return of the inositol phosphates. Nat Rev Mol Cell Biol 2: 327–338

40. Kim S, Bhandari R, Brearley CA, Saiardi A (2024) The inositol phosphate signalling network in physiology and disease. Trends Biochem Sci 49: 969–985

41. Kraszewska E (2008) The plant Nudix hydrolase family. Acta Biochimica Polonica 55: 663–671

42. Kurtović K, Schmidt V, Nehasilová M, Vosolsobě S, Petrášek J (2023) Rediscovering *Chara* as a model organism for molecular and evo-devo studies. Protoplasma. doi: 10.1007/s00709-023-01900-3

43. Laha D, Johnen P, Azevedo C, Dynowski M, Weiß M, Capolicchio S, Mao H, Iven T, Steenbergen M, Freyer M, et al (2015) VIH2 regulates the synthesis of inositol pyrophosphate InsP8 and jasmonate-dependent defenses in *Arabidopsis*. Plant Cell 27: 1082–1097

44. Laha D, Kamleitner M, Johnen P, Schaaf G (2021a) Analyses of inositol phosphates and Phosphoinositides by strong anion exchange (SAX)-HPLC. In D Bartels, P Dörmann, eds, Plant Lipids: Methods and Protocols. Springer US, New York, NY, pp 365–378

45. Laha D, Parvin N, Dynowski M, Johnen P, Mao H, Bitters ST, Zheng N, Schaaf G (2016) Inositol polyphosphate binding specificity of the jasmonate receptor complex. Plant Physiol 171: 2364–2370

46. Laha D, Parvin N, Hofer A, Giehl RFH, Fernandez-Rebollo N, von Wirén N, Saiardi A, Jessen HJ, Schaaf G (2019) Arabidopsis ITPK1 and ITPK2 have an evolutionarily conserved phytic acid kinase activity. ACS Chem Biol 14: 2127–2133

47. Laha D, Portela-Torres P, Desfougères Y, Saiardi A (2021b) Inositol phosphate kinases in the eukaryote landscape. Advances in Biological Regulation 79: 100782

48. Lampe S, Mohanty TK, Bhandari R, Fiedler D (2025) Protein pyrophosphorylation by inositol pyrophosphates - detection, function, and regulation. FEBS Lett. doi: 10.1002/1873-3468.70240

49. Laurent F, Bartsch SM, Shukla A, Rico-Resendiz F, Couto D, Fuchs C, Nicolet J, Loubéry S, Jessen HJ, Fiedler D, et al (2024) Inositol pyrophosphate catabolism by three families of phosphatases regulates plant growth and development. PLOS Genetics 20: e1011468

50. Leebens-Mack JH, Barker MS, Carpenter EJ, Deyholos MK, Gitzendanner MA, Graham SW, Grosse I, Li Z, Melkonian M, Mirarab S, et al (2019) One thousand plant transcriptomes and the phylogenomics of green plants. Nature 574: 679–685

51. Liu G, Dürr-Mayer T, Lu M, Jessen HJ (2025) Establishing 18O-labeled inositol phosphates for quantitative capillary electrophoresis-mass spectrometry: fragmentation pathways and comparison with 13C-labeled analogs. Anal Chem 97: 25282–25294

52. Liu G, Riemer E, Schneider R, Cabuzu D, Bonny O, Wagner CA, Qiu D, Saiardi A, Strauss A, Lahaye T, et al (2023) The phytase RipBL1 enables the assignment of a specific inositol phosphate isomer as a structural component of human kidney stones. RSC Chem Biol 4: 300–309

53. Lorenzo-Orts L, Couto D, Hothorn M (2020) Identity and functions of inorganic and inositol polyphosphates in plants. New Phytologist 225: 637–652

54. Martin WF, Allen JF (2018) An Algal Greening of Land. Cell 174: 256–258

55. McCourt RM, Karol KG, Guerlesquin M, Feist M (1996) Phylogeny of extant genera in the family Characeae (Charales, Charophyceae) based on rbcL, sequences and morphology. American Journal of Botany 83: 125–131

56. McCourt RM, Karol KG, Hall JD, Casanova MT, Grant MC (2017) Charophyceae (Charales). *In* JM Archibald, AGB Simpson, CH Slamovits, eds, Handbook of the Protists. Springer International Publishing, Cham, pp 165–183

57. Meng EC, Goddard TD, Pettersen EF, Couch GS, Pearson ZJ, Morris JH, Ferrin TE (2023) UCSF ChimeraX: tools for structure building and analysis. Protein Sci e4792

58. Miller GJ, Wilson MP, Majerus PW, Hurley JH (2005) Specificity determinants in inositol polyphosphate synthesis: crystal structure of inositol 1,3,4-trisphosphate 5/6-kinase. Molecular Cell 18: 201–212

59. Morales-Pineda M, García-Gómez ME, Bedera-García R, García-González M, Couso I (2023) CO2 levels modulate carbon utilization, energy levels and inositol polyphosphate profile in chlorella. Plants 12: 129

60. Mulugu S, Bai W, Fridy PC, Bastidas RJ, Otto JC, Dollins DE, Haystead TA, Ribeiro AA, York JD (2007) A conserved family of enzymes that phosphorylate inositol hexakisphosphate. Science 316: 106–109

61. Nagpal L, He S, Rao F, Snyder SH (2024) Inositol pyrophosphates as versatile metabolic messengers. Annual Review of Biochemistry 93: 317–338

62. Nishiyama T, Sakayama H, de Vries J, Buschmann H, Saint-Marcoux D, Ullrich KK, Haas FB, Vanderstraeten L, Becker D, Lang D, et al (2018) The *Chara* genome: secondary complexity and implications for plant terrestrialization. Cell 174: 448–464.e24

63. Notredame C, Higgins DG, Heringa J (2000) T-coffee: a novel method for fast and accurate multiple sequence alignment1. Journal of Molecular Biology 302: 205–217

64. Otterhag L, Sommarin M, Pical C (2001) N-terminal EF-hand-like domain is required for phosphoinositide-specific phospholipase C activity in *Arabidopsis thaliana*. FEBS Letters 497: 165–170

65. Plevin MJ, Mills MM, Ikura M (2005) The LxxLL motif: a multifunctional binding sequence in transcriptional regulation. Trends Biochem Sci 30: 66–69

66. Pullagurla NJ, Shome S, Liu G, Jessen HJ, Laha D (2025) Orchestration of phosphate homeostasis by the ITPK1-type inositol phosphate kinase in the liverwort *Marchantia polymorpha*. Plant Physiol 197: kiae454

67. Qiu D, Gu C, Liu G, Ritter K, B. Eisenbeis V, Bittner T, Gruzdev A, Seidel L, Bengsch B, B. Shears S, et al (2023) Capillary electrophoresis mass spectrometry identifies new isomers of inositol pyrophosphates in mammalian tissues. Chemical Science 14: 658–667

68. Qiu D, Wilson MS, Eisenbeis VB, Harmel RK, Riemer E, Haas TM, Wittwer C, Jork N, Gu C, Shears SB, et al (2020) Analysis of inositol phosphate metabolism by capillary electrophoresis electrospray ionization mass spectrometry. Nat Commun 11: 6035

69. Raia P, Lee K, Bartsch SM, Rico-Resendiz F, Portugal-Calisto D, Vadas O, Panse VG, Fiedler D, Hothorn M (2025) A small signaling domain controls PPIP5K phosphatase activity in phosphate homeostasis. Nat Commun 16: 1753

70. Rambaut A (2018) FigTree software, version 1.4.4. http://tree.bio.ed.ac.uk/software/figtree/

71. Rambaut A, Drummond AJ, Xie D, Baele G, Suchard MA (2018) Posterior summarization in bayesian phylogenetics using tracer 1.7. Systematic Biology 67: 901–904

72. Rana P, Edathil Kadangodan A, Koley P, Ghosh R, Pullagurla NJ, Naik HC, Mondal P, Gayen S, Laha D (2025) GA-independent DELLA regulation by inositol pyrophosphate in a nonvascular land plant. Nat Chem Biol 21: 1697–1708

73. Randall TA, Gu C, Li X, Wang H, Shears SB (2020) A two-way switch for inositol pyrophosphate signaling: Evolutionary history and biological significance of a unique, bifunctional kinase/phosphatase. Advances in Biological Regulation 75: 100674

74. Riemer E, Pullagurla NJ, Yadav R, Rana P, Jessen HJ, Kamleitner M, Schaaf G, Laha D (2022) Regulation of plant biotic interactions and abiotic stress responses by inositol polyphosphates. Frontiers in Plant Science 13:

75. Riemer E, Qiu D, Laha D, Harmel RK, Gaugler P, Gaugler V, Frei M, Hajirezaei M-R, Laha NP, Krusenbaum L, et al (2021) ITPK1 is an InsP6/ADP phosphotransferase that controls phosphate signaling in *Arabidopsis*. Mol Plant 14: 1864–1880

76. Ritter K, Braun A-SC, Liu G, Lu M, Gaugler V, Schaaf G, Jessen HJ (2025) Expanding the inositol pyrophosphate toolbox: stereoselective synthesis and application of PP-InsP4 isomers in plant signaling. Angew Chem Int Ed Engl 64: e202507058

77. Ritter K, Gaugler V, Stolze SC, Ghosh R, Jayamon A, Yadav R, Wollensack F, Laha D, Nakagami H, Schaaf G, et al (2026) Affinity-based interactome mapping of inositol pyrophosphates reveals 4/6-PP-InsP5-binding proteins in plants. Adv Sci (Weinh) e24290

78. Sanz-Luque E, Bhaya D, Grossman AR (2020) Polyphosphate: a multifunctional metabolite in cyanobacteria and algae. Front Plant Sci. doi: 10.3389/fpls.2020.00938

79. Schneider R, Lami K, Prucker I, Stolze SC, Strauß A, Schmidt JM, Bartsch SM, Langenbach K, Lange E, Ritter K, et al (2025) NUDIX hydrolases target specific inositol pyrophosphates and regulate phosphate homeostasis and bacterial pathogen susceptibility in *Arabidopsis*. J Integr Plant Biol 67: 3123–3151

80. Shears SB (2015) Inositol pyrophosphates: why so many phosphates? Adv Biol Regul 57: 203–216

81. Shears SB, Wang H (2019) Inositol phosphate kinases: Expanding the biological significance of the universal core of the protein kinase fold. Advances in Biological Regulation 71: 118–127

82. Shukla A, Kaur M, Kanwar S, Kaur G, Sharma S, Ganguli S, Kumari V, Mazumder K, Pandey P, Rouached H, et al (2021) Wheat inositol pyrophosphate kinase TaVIH2-3B modulates cell-wall composition and drought tolerance in *Arabidopsis*. BMC Biol 19: 261

83. Sturm K, Pri-Tal O, Rico-Resendiz F, Verma Y, Richter A, Chen H, Broger L, Hothorn LA, Fiedler D, Panse VG, et al (2025) An inositol pyrophosphate interaction screen provides insight into the regulation of plant casein kinase II. 2025.11.27.691040

84. Tasma IM, Brendel V, Whitham SA, Bhattacharyya MK (2008) Expression and evolution of the phosphoinositide-specific phospholipase C gene family in *Arabidopsis thaliana*. Plant Physiology and Biochemistry 46: 627–637

85. The Inkscape Developers (2024) Inkscape - Draw Freely. | Inkscape.

86. The UniProt Consortium (2025) UniProt: the Universal Protein Knowledgebase in 2025. Nucleic Acids Research 53: D609–D617

87. Wickett NJ, Mirarab S, Nguyen N, Warnow T, Carpenter E, Matasci N, Ayyampalayam S, Barker MS, Burleigh JG, Gitzendanner MA, et al (2014) Phylotranscriptomic analysis of the origin and early diversification of land plants. PNAS 111: E4859–E4868

88. Wild R, Gerasimaite R, Jung J-Y, Truffault V, Pavlovic I, Schmidt A, Saiardi A, Jessen HJ, Poirier Y, Hothorn M, et al (2016) Control of eukaryotic phosphate homeostasis by inositol polyphosphate sensor domains. Science 352: 986–990

89. Williams SP, Gillaspy GE, Perera IY (2015) Biosynthesis and possible functions of inositol pyrophosphates in plants. Frontiers in Plant Science 6:

90. Wilson MS, Saiardi A (2018) Inositol phosphates purification using titanium dioxide beads. Bio Protoc 8: e2959

91. Xiong T, Zhang Z, Fan T, Ye F, Ye Z (2024) Origin, evolution, and diversification of inositol 1,4,5-trisphosphate 3-kinases in plants and animals. BMC Genomics 25: 350

92. Yadav R, Liu G, Rana P, Pullagurla NJ, Qiu D, Jessen HJ, Laha D (2025) Conservation of heat stress acclimation by the IPK2-type kinases that control the synthesis of the inositol pyrophosphate 4/6-InsP7 in land plants. PLOS Genetics 21: e1011838

93. Yang S-Y, Lin W-Y, Hsiao Y-M, Chiou T-J (2024) Milestones in understanding transport, sensing, and signaling of the plant nutrient phosphorus. Plant Cell 36: 1504–1523

94. Yoshimura K, Shigeoka S (2015) Versatile physiological functions of the Nudix hydrolase family in Arabidopsis. Biosci Biotechnol Biochem 79: 354–366

95. Zhang Y, Skolnick J (2004) Scoring function for automated assessment of protein structure template quality. Proteins: Structure, Function, and Bioinformatics 57: 702–710

96. Zhu J, Lau K, Puschmann R, Harmel RK, Zhang Y, Pries V, Gaugler P, Broger L, Dutta AK, Jessen HJ, et al (2019) Two bifunctional inositol pyrophosphate kinases/phosphatases control plant phosphate homeostasis. eLife 8: e43582

