## Supplemental Figures and Tables for "Inositol phosphates, pyrophosphates and the genes involved in their turnover in the streptophyte green alga *Chara braunii*"

*Co-sharing first authors

[#]Co-corresponding authors

**Supplementary Information**

**Supplementary Tables:** p. 2

**Supplementary Figures:** p. 6

**Supplementary References:** p. 17

### Supplementary Tables

**Table S1. Complete list of *Chara braunii* homologs of *A. thaliana* proteins involved in InsP and PP-InsP production*.*** Putative proteins with sequence and structural similarity to selected, known plant enzymes involved in InsP metabolism are listed. The accession number of the respective query protein in *A. thaliana* is given, followed by the enzyme name, respective protein domains, the accession number of the best-matching protein in *C. braunii*, followed by the percentage of identical amino acids and the length of the sequence overlap (including gaps). Putative InsP enzymes from *C. braunii* were identified by sequence similarity using BLASTP (Altschul et al., 1990), or by similarities in domain architecture using INTERPRO (Blum et al., 2025). Note that neither the biochemical activities, nor the sequence identities *in vivo* have been experimentally assessed for the listed *Chara* proteins. BLASTP results below 30% query cover or alignment e-values above 1e-5 have been omitted. This list complements **Table 1**.

| **Accession no.** | **Enzyme** | **Domain** | **Chara homolog** | **Sequence identity** | **Overlap** |
|---|---|---|---|---|---|
| **Kinases mediating the synthesis of InsPs** | | | | | |
| NP_196354 | IPK2a | IPK_sf | g44488* | 39% | 286/205 |
| | | IPK_sf | g16917 | 34% | 286/514 |
| NP_001331991 | IPK2b | IPK_sf | g44488* | 39% | 300/205 |
| | | IPK_sf | g16917 | 36% | 300/514 |
| Q93YN9 | IPK1 | IP5_2-K_N_lobe | g22893 | 33% | 451/1350 |
| Q9SBA5 | ITPK1 | ITPK1_N | g23313* | 44% | 319/340 |
| | | InsP(3)kin | g54337* | 32% | 319/325 |
| | | Ins134_P3_kin | g3607 | 27% | 319/638 |
| O81893 | ITPK2 | ITPK1_N | g23313* | 56% | 391/340 |
| | | InsP(3)kin | g54337* | 36% | 391/325 |
| | | Ins134_P3_kin | g3607 | 25% | 391/638 |
| F4J8C6 | VIH1 | His_PPase superfam | g24415* | 68% | 1049/1524 |
| Q84WW3 | VIH2 | His_PPase superfam | g24415* | 68% | 1050/1524 |
| **Phosphoinositide-specific phospholipases** | | | | | |
| Q39033 | PLC2 | PI-PLC Y-box | g36522* | 44% | 581/1860 |
| | | | g23178 | 50% | 581/2377 |
| | | | g24440 | 70% | 581/711 |
| Q9LY51 | PLC7 | PI-PLC Y-box | g36522* | 43% | 584/1860 |
| | | | g24440 | 31% | 584/711 |
| | | | g23178 | 40% | 584/2377 |
| Q944C1 | PLC4 | PI-PLC Y-box | g36522 | 43% | 597/1860 |
| | | | g24440 | 62% | 597/711 |
| **Phosphohydrolases** | | | | | |

| | | | | | |
|---|---|---|---|---|---|
| Q9ZVN4 | DSP1 | Prot-tyrosine_ phosphatase-like | g23885 | 69% | 146/215 |
| | | | g29374* | 41% | 406/215 |
| | | | g30804* | 32% | 235/215 |
| | | | g19745 | 37% | 294/215 |
| Q84MD6 | DSP2 | Prot-tyrosine_ phosphatase-like | g23885 | 54% | 146/257 |
| | | | g29374* | 33% | 406/257 |
| | | | g19745* | 39% | 294/257 |
| | | | g30804 | 29% | 235/257 |
| | | | g23883 | 39% | 128/257 |
| Q681Z2 | DSP3 | Prot-tyrosine_ phosphatase-like | g23885 | 51% | 146/203 |
| | | | g29374* | 39% | 406/203 |
| | | | g30804* | 36% | 235/203 |
| | | | g19745 | 38% | 294/203 |
| | | | g23883 | 45% | 128/203 |
| Q940L5 | DSP4 | Prot-tyrosine_ phosphatase-like | g23885 | 66% | 146/198 |
| | | | g29374* | 41% | 406/198 |
| | | | g19745* | 43% | 294/198 |
| | | | g30804 | 35% | 235/198 |
| | | | g23883 | 44% | 128/198 |
| Q9FFD7 | DSP5 | Prot-tyrosine_ phosphatase-like | g23885 | 57% | 146/204 |
| | | | g29374* | 36% | 406/204 |
| | | | g30804* | 36% | 235/204 |
| | | | g19745 | 44% | 294/204 |
| | | | g23883 | 47% | 294/204 |
| Q9LE73 | NUDT4 | NUDIX_hydrolase _CS | g45972* | 35% | 566/207 |
| Q9ZU95 | NUDT17 | NUDIX_hydrolase _CS | g45972* | 37% | 566/182 |
| Q9LQU5 | NUDT18 | NUDIX_hydrolase _CS | g45972* | 38% | 566/176 |
| Q8VY81 | NUDT21 | NUDIX_hydrolase _CS | g45972* | 35% | 566/198 |
| Q93ZY7 | NUDT12 | NUDIX_hydrolase _CS | g45972* | 35% | 566/203 |
| Q52K88 | NUDT13 | NUDIX_hydrolase _CS | g45972* | 41% | 566/202 |
| Q9LHK1 | NUDT16 | NUDIX_hydrolase _CS | g45972* | 42% | 566/180 |

**Table S2. Instrument and scan source parameters of QQQ.**

| Parameter | Setting |
|---|---|
| Gas temperature | 150 °C |
| Gas flow | 11 L $min^{-1}$ |
| Nebulizer pressure | 8 psi |
| Sheath gas temperature | 175 °C |
| Sheath gas flow | 8 L $min^{-1}$ |
| Capillary voltage | −2000 V |
| Nozzle voltage | 2000 V |
| High-pressure RF (ion funnel) | 70 V |
| Low-pressure RF (ion funnel) | 40 V |

**Table S3. MRM transitions setting of InsPs for *Chara* samples.**

| Molecular name | Precursor ion (m/z) | Product ion (m/z) | Dwell (ms) | Fragmentor (V) | Collision energy (V) | Cell accelerator (V) | Polarity |
|---|---|---|---|---|---|---|---|
| $InsP_3$ | 418.9 | 320.8 | 50 | 166 | 17 | 4 | Negative |
| $^{13}C_6$ -$InsP_3$ | 424.9 | 326.8 | 50 | 166 | 17 | 4 | Negative |
| $InsP_4$ | 499.0 | 418.9 | 50 | 166 | 5 | 1 | Negative |
| $^{13}C_6$ -$InsP_4$ | 505.0 | 424.9 | 50 | 166 | 5 | 1 | Negative |
| $InsP_5$ | 579.0 | 498.9 | 50 | 166 | 9 | 3 | Negative |
| $^{13}C_6$-$InsP_5$ | 585.0 | 504.9 | 50 | 166 | 9 | 3 | Negative |
| $InsP_6$ | 659.0 | 480.9 | 50 | 166 | 13 | 4 | Negative |
| $^{13}C_6$-$InsP_6$ | 665.0 | 486.9 | 50 | 166 | 13 | 4 | Negative |
| PP-$InsP_5$ | 739.0 | 319.9 | 50 | 166 | 9 | 3 | Negative |
| $^{13}C_6$- 5-PP-InsP5 | 745.0 | 322.9 | 50 | 166 | 9 | 3 | Negative |
| $^{18}O_6$-4- PP-$InsP_5$ | 751.0 | 327.9 | 50 | 166 | 9 | 3 | Negative |
| $^{13}C_6$-$(PP)_2$-$InsP_4$ | 825.0 | 362.8 | 50 | 166 | 9 | 1 | Negative |

**Table S4. Isotopic internal standards and final working concentrations for quantitative CE–MS analysis**

| **Isotopic internal standard** | **Inositol phosphate species** | **Final concentration in sample (μM)** |
|---|---|---|
| $^{13}C_6$-Ins(1,2,3)$P_3$ | Inositol 1,2,3-trisphosphate | 10 |
| $^{13}C_6$-Ins(1,3,4,5,6)$P_5$ | Inositol pentakisphosphate (2-OH-Ins$P_5$) | 4 |
| $^{13}C_6$-Ins$P_6$ | Inositol hexakisphosphate (Ins$P_6$) | 8 |
| $^{13}C_6$-5-PP-Ins$P_5$ | 5-diphospho-inositol pentakisphosphate | 1 |
| $^{13}C_6$-1- PP-Ins$P_5$ | 1-diphospho-inositol pentakisphosphate | 1 |
| $^{13}C_6$-1,5-Ins$P_8$ | 1,5-bis-diphospho-inositol tetrakisphosphate | 0.5 |
| $^{18}O_2$-4- PP-Ins$P_5$ | 4-diphospho-inositol pentakisphosphate | 1 |

### Supplementary Figures

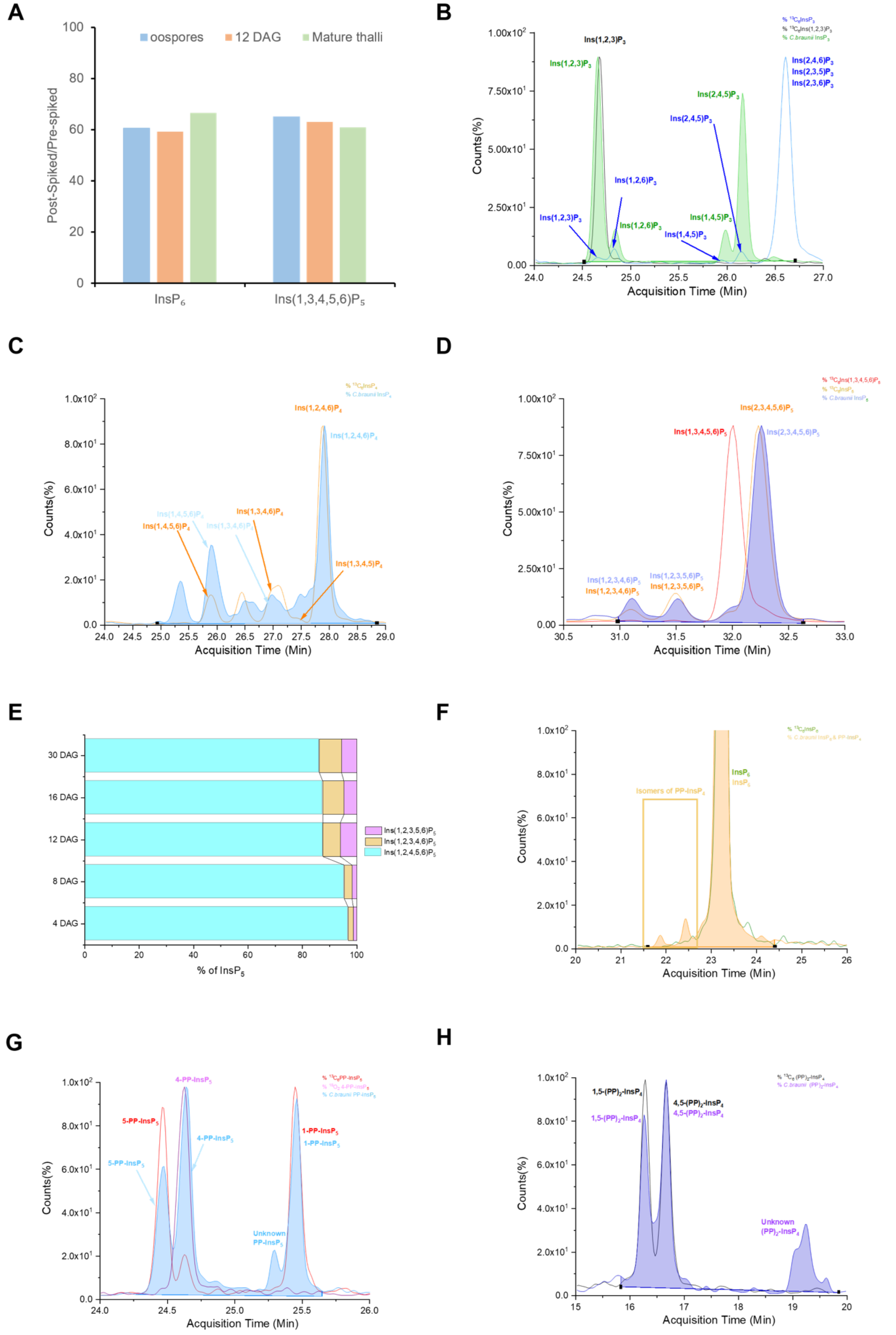

**Figure S1. Isomer-resolved CE–ESI–MS profiling of InsPs and PP-InsPs in *Chara braunii*. A)** Relative abundance of $InsP_6$ and Ins(1,3,4,5,6)$P_5$ across developmental stages (oospores, 12 DAG, mature thalli), shown as percentage of total signal. **B)** Representative extracted-ion electropherograms showing separation of $InsP_3$ isomers, including Ins(1,2,3)$P_3$, Ins(1,2,6)$P_3$, Ins(1,4,5)$P_3$, and Ins(2,4,5)$P_3$. **C)** Extracted-ion electropherograms of $InsP_4$ isomers. Ins(1,4,5,6)$P_4$ and the co-migrating pair Ins(1,2,4,6)$P_4$ / Ins(2,3,4,6)$P_4$ represent the dominant species. **D)** Extracted-ion electropherograms of $InsP_5$ isomers showing multiple resolved species, with the predominant signal corresponding to Ins(1,2,4,5,6)$P_5$ / Ins(2,3,4,5,6)$P_5$. **E)** Relative distribution of $InsP_5$ isomers across developmental stages. **F)** $InsP_6$ electropherograms showing a single dominant peak of $InsP_6$ and two isomer of PP-$InsP_4$. **G)** Separation of PP-$InsP_5$ isomers, including 1-PP-$InsP_5$, 4-PP-$InsP_5$, and 5-PP-$InsP_5$. **H)** Detection of $(PP)_2$-$InsP_4$ species with additional unassigned isomers present. Isomer nomenclature follows the lowest-numbering convention; unresolved enantiomeric pairs are reported with subscript numbering.

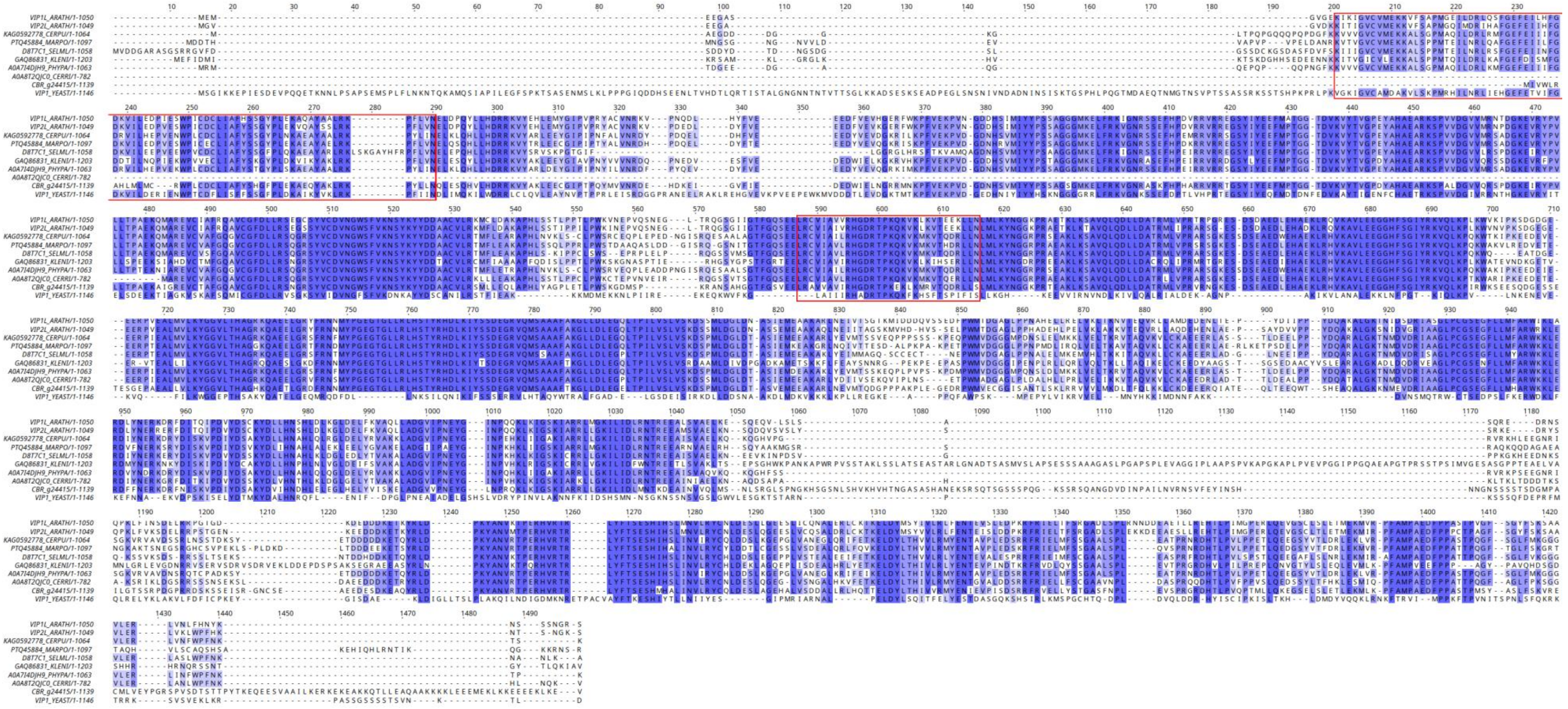


**Figure S2. Sequence alignment of VIP candidates from *Chara braunii* with representative sequences from viridiplantae and yeast.** Sequence comparison of inositol hexakisphosphate and diphosphoinositol-pentakisphosphate kinase homologs from *Arabidopsis thaliana, Ceratodon purpureus*, *Marchantia polymorpha*, *Selaginella moellendorffii*, *Klebsormidium nitens, Physcomitrium patens, Chara braunii* and *Ceratopteris richardii* plus the fungal model *Saccharomyces cerevisiae* VIP1. The multiple sequence alignment was generated using JalView v.2.11.2.7 (default colored percentage identity). Conserved diphosphoinositol pentakisphosphate kinase 2 N-terminal domain and histidine acid phosphatases phosphohistidine signature are boxed in red. This figure extends **Fig. 5**.

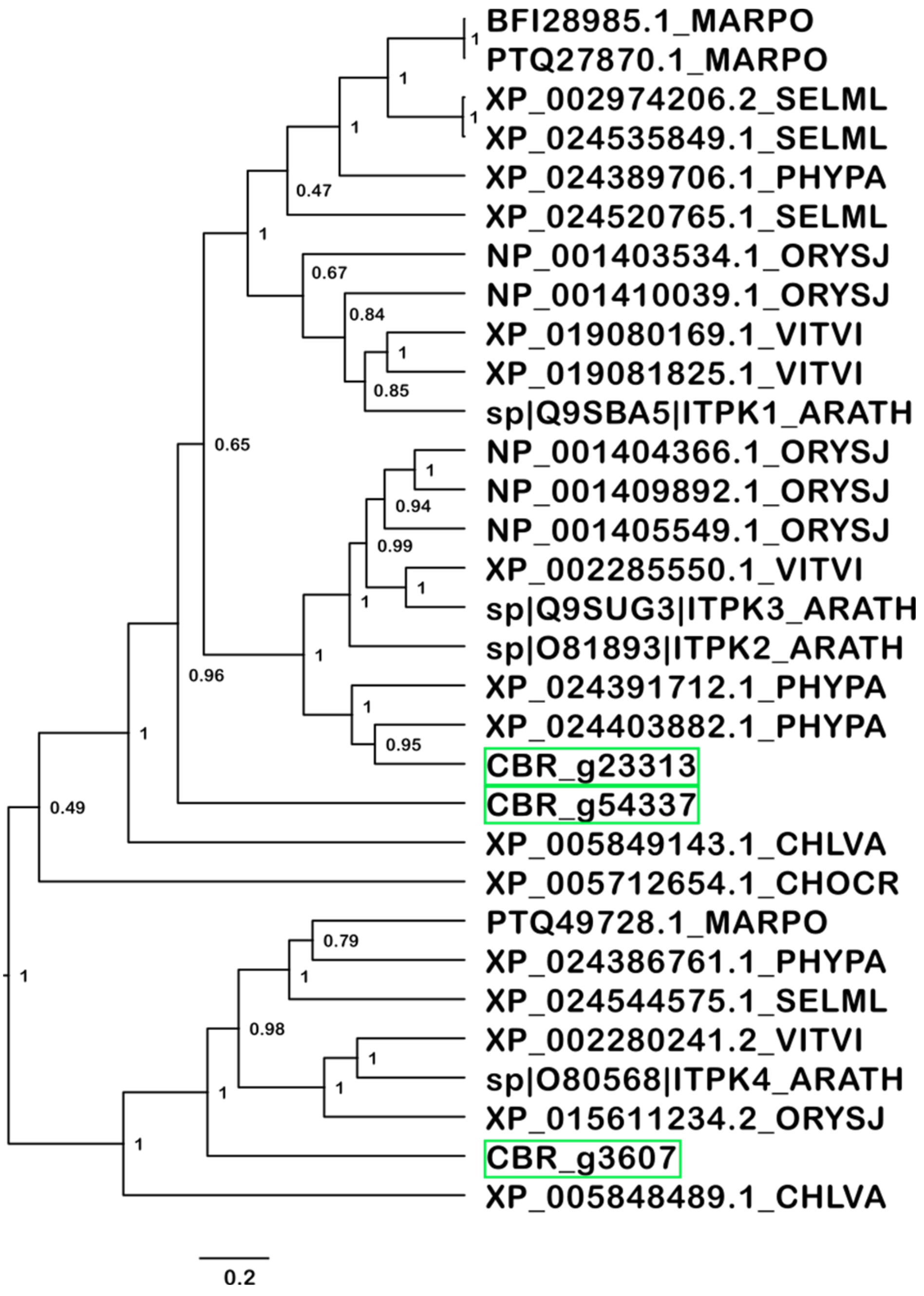


**Figure S3. Phylogenetic relationships of *C. braunii* ITPK candidates.** Bayesian inference of three putative *C. braunii* ITPK homologs (boxed in green). The dataset of sequences for alignment (see **Fig. S4**) and phylogenetic tree generation was based on previous work (Laha et al., 2019; Pullagurla et al., 2025).

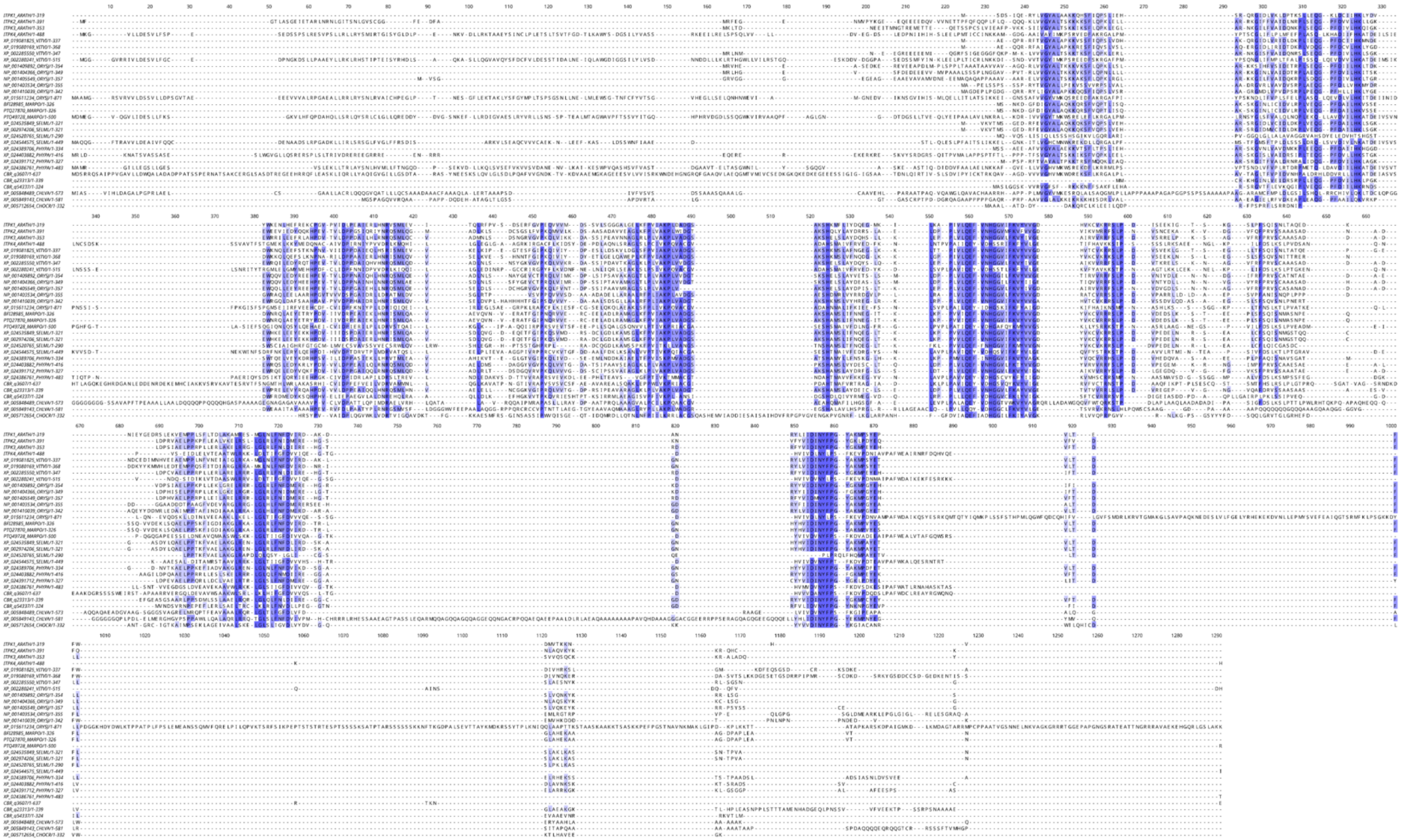

**Figure S4. Sequence alignment of ITPK candidates in *Chara braunii*.** Complete alignment of ITPK homolog amino acid sequences from selected viridiplantae model organisms *Arabidopsis thaliana*, *Vitis vinifera*, *Oryza sativa* subsp. *japonica, Marchantia polymorpha, Physcomitrium patens,*

*Selaginella moellendorffii, Chara braunii, Chlorella variabilis* and *Chondrus crispus* was generated using JalView v.2.11.2.7 (default colored percentage identity). This figure extends **Fig. S3**.

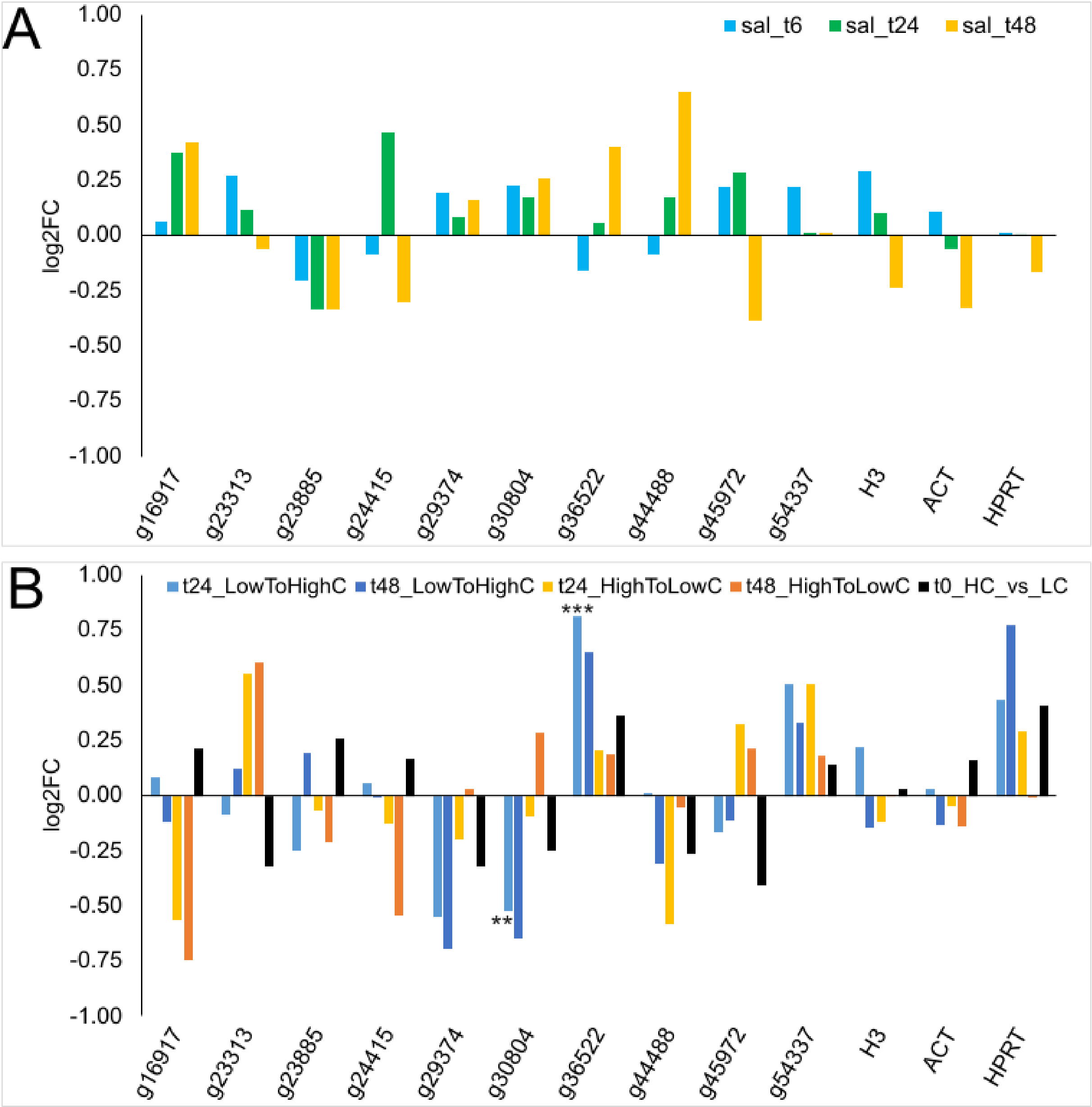


**Figure S5. Differential expression of selected InsP gene candidates.** $Log_2$ fold changes were plotted for selected InsP metabolism gene candidates of interest utilizing data from two previously published *Chara braunii* transcriptome analyses. **A)** Impact of 5 PSU salt stress on the gene expression. **B)** Gene expression changes in *Chara braunii* under shifting inorganic carbon (Ci) conditions (low to high Ci, high to low Ci, high compared to low Ci). Asterisks indicate significant changes within the context of the examined publications compared to (i) t0 normal growth and (ii) t0 acclimated low/ high carbon treatment (***, padj ≤0.01; ** padj ≤0.05). Gene expression data were collected from Heß et al. (2023a) and Heise et al. (2025)

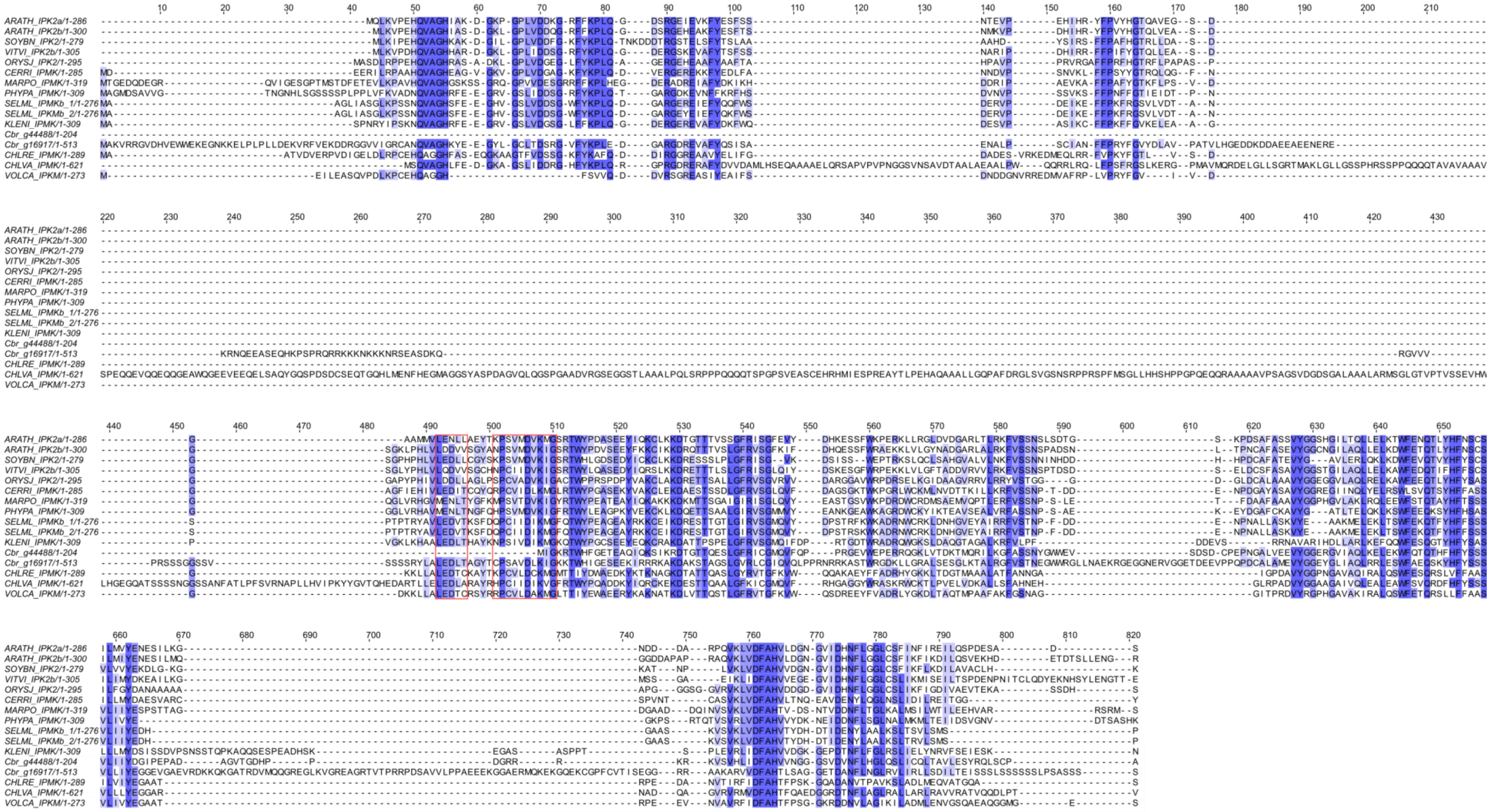

**Figure S6. Sequence alignment of IPMK candidates in *Chara braunii*.** Complete alignment of IPMK central sequences from AtIPK2a/b homologs in viridiplantae model organisms *Glycine max, Vitis vinifera, Oryza sativa* subsp. *japonica, Ceratopteris richardii, Marchantia polymorpha, Physcomitrium patens, Selaginella moellendorffii, Klebsormidium nitens, Chara braunii, Chlamydomonas reinhardtii, Chlorella variabilis and Volvox*

*carteri*. Multiple sequence alignment was generated using M-coffee and JalView v.2.11.2.7 (default colored percentage identity). Conserved LxxLL protein recognition and PxxxDxKxG catalytic motifs are highlighted in red squares. This figure extends **Fig. 4**.

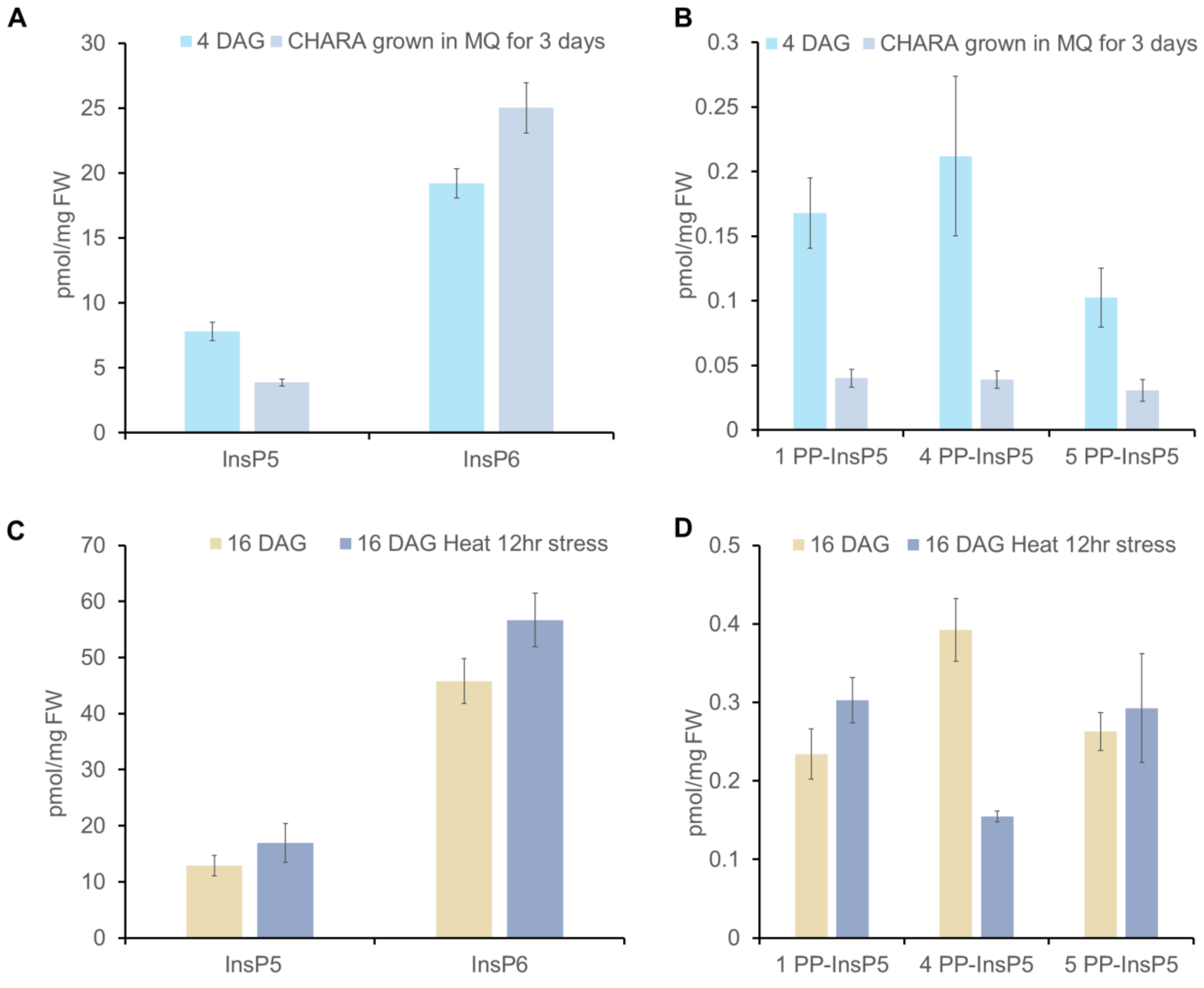


**Figure S7. Environmental modulation of inositol phosphate metabolism in *Chara braunii.*** **A)** Quantification of total $InsP_5$ and $InsP_6$ in 4 DAG *Chara braunii* compared with samples grown in MQ water for 3 days. **B)** Quantification of PP-$InsP_5$ isomers in 4 DAG *Chara braunii* and MQ-treated samples. All three PP-$InsP_5$ isomers (1-PP-$InsP_5$, 4-PP-$InsP_5$, and 5-PP-$InsP_5$) are reduced in MQ-treated samples compared with untreated 4 DAG controls. **C)** Quantification of total $InsP_5$ and $InsP_6$ in 16 DAG samples under heat stress (12 h) compared with untreated 16 DAG controls. **D)** Quantification of PP-$InsP_5$ isomers in 16 DAG samples following heat stress (12 h). 1-PP-$InsP_5$ and 5-PP-$InsP_5$ show moderate increases, whereas 4-PP-$InsP_5$ decreases relative to untreated controls. Data represent mean ± SD; n = 3 biological replicates.

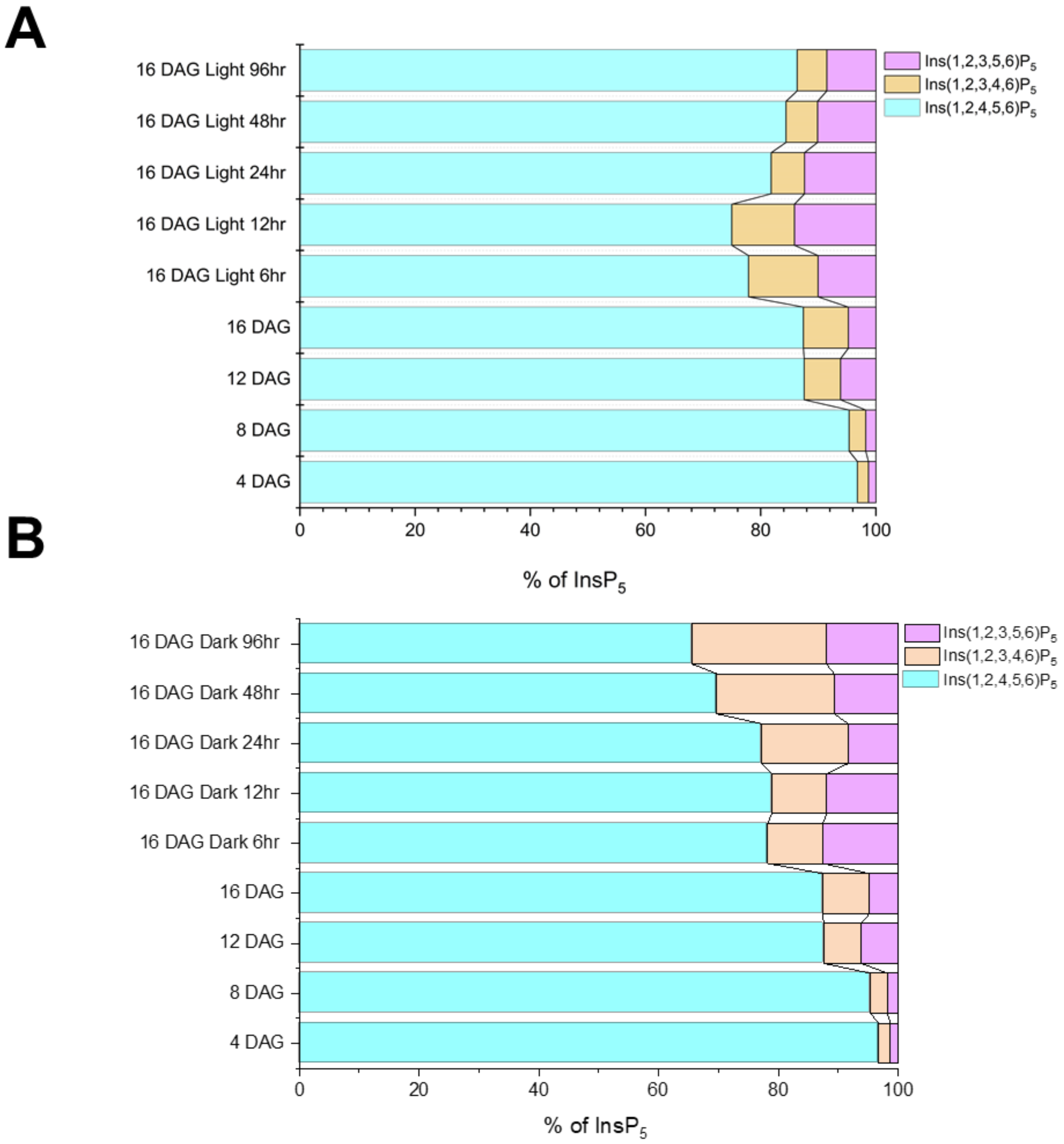


**Figure S8 Relative distribution of $InsP_5$ isomers in response to light and dark exposure in *Chara braunii*. A)** Stacked bar representation showing the percentage of $InsP_5$ isomers during extended light exposure. **B)** Stacked bar representation showing the percentage of $InsP_5$ isomers during extended dark exposure.

**Supplementary References**


**Altschul SF, Gish W, Miller W, Myers EW, Lipman DJ** (1990) Basic local alignment search tool. J Mol Biol **215**: 403–410

**Blum M, Andreeva A, Florentino LC, Chuguransky SR, Grego T, Hobbs E, Pinto BL, Orr A, Paysan-Lafosse T, Ponamareva I, et al** (2025) InterPro: the protein sequence classification resource in 2025. Nucleic Acids Research **53**: D444–D456

**Heise CM, Heß DA, Walke P, Voß M, Schubert H, Hess WR, Hagemann M** (2025) Evidence for a $CO_2$-concentrating mechanism in the model streptophyte green alga *Chara braunii*. New Phytologist **247**: 1218–1233

**Heß D, Heise CM, Schubert H, Hess WR, Hagemann M** (2023) The impact of salt stress on the physiology and the transcriptome of the model streptophyte green alga *Chara braunii*. Physiologia Plantarum **175**: e14123

**Laha D, Parvin N, Hofer A, Giehl RFH, Fernandez-Rebollo N, von Wirén N, Saiardi A, Jessen HJ, Schaaf G** (2019) *Arabidopsis* ITPK1 and ITPK2 have an evolutionarily conserved phytic acid kinase activity. ACS Chem Biol **14**: 2127–2133

**Pullagurla NJ, Shome S, Liu G, Jessen HJ, Laha D** (2025) Orchestration of phosphate homeostasis by the ITPK1-type inositol phosphate kinase in the liverwort *Marchantia polymorpha*. Plant Physiol **197**: kiae454
